# Re-evaluating homoploid reticulate evolution in the annual sunflowers

**DOI:** 10.1101/2022.10.14.512273

**Authors:** Gregory L. Owens, Kaichi Huang, Marco Todesco, Loren H. Rieseberg

## Abstract

Sunflowers of the genus *Helianthus* are models for hybridization research and contain three of the best studied examples of homoploid hybrid speciation. To understand the broader picture of hybridization within the annual sunflowers, we used whole genome resequencing to conduct a phylogenomic analysis and test for gene flow between lineages. We find that all annual sunflower species tested have evidence of admixture, suggesting hybridization was common during the radiation of the genus. Support for the major species tree decreases with recombination rate, consistent with hybridization and introgression contributing to discordant topologies. Admixture graphs found hybridization to be associated with the origins of the three putative hybrid species (*H. anomalus, H. deserticola*, and *H. paradoxus*). However, the hybridization events are more ancient than suggested by previous work. Furthermore, *H. anomalus* and *H. deserticola* appear to have arisen from a single hybridization event involving an unexpected donor, rather than through multiple independent events as previously proposed. Using a broader data set that covers the whole *Helianthus* genus, including perennial species, we find that signals of introgression span the genus and beyond, suggesting highly divergent introgression and/or the sorting of ancient haplotypes. Thus, *Helianthus* can be viewed as a syngameon in which largely reproductively isolated species are linked together by occasional or frequent gene flow.

## Introduction

The evolutionary importance of hybridization and introgression has been explored since Darwin’s time (Darwin 1859). While initially zoologists thought that hybridization was relatively rare and unimportant (Mayr 1963), botanists have long emphasized the role of hybridization and introgression in both adaptation and speciation (Anderson 1949; Heiser 1949; Anderson and Stebbins 1954). Genomics eventually proved botanists correct in this regard; evidence of past hybridization is common in both animals and plants (Figueiró et al. 2017; Fontaine et al. 2015; Mallet et al. 2016; Calfee et al. 2021; Dagilis et al. 2021). In several cases, alleles responsible for ecologically important traits have been found to be acquired through introgression (Suarez-Gonzalez et al. 2018; Jones et al. 2018; Oziolor et al. 2019), supporting the notion that introgression not only occurs, but can be evolutionarily important.

Hybridization can also result in the formation of new species through homoploid hybrid speciation, in which an admixed lineage acquires reproductive isolation from its progenitors through a combination of alleles or chromosomal rearrangements from both parents (Grant 1958; Schumer et al. 2014; Stebbins 1957). One way this can occur is if two parental species differ at both habitat and mate choice. A hybrid population between these two may have alternate parental alleles for these two traits, and therefore be separated from one parental species by habitat and the other by mate choice (Wang et al., 2021). Hybrid speciation can also occur when hybridization results in the ability to colonize a new ecological niche, reject parental species as potential mates or have unique combinations of chromosomal rearrangements (Gross and Rieseberg 2005; Melo et al. 2009; Rieseberg et al. 1995). Schumer et al. (2014) argued that three criteria must be satisfied to demonstrate homoploid hybrid speciation: (1) reproductive isolation from parental species, (2) prior admixture, and (3) evidence that reproductive barriers arose via hybridization. Evaluation of the natural hybridization literature indicates that while admixed lineages are relatively common (criterion 2), few case studies satisfy criteria 1 and 3 (Schumer et al. 2014). Three examples of hybrid species thought to address all three criteria are within the sunflower genus, *Helianthus*.

The widespread species *Helianthus annuus* and *H. petiolaris* are thought to be the parents of three independent homoploid hybrid species, *H. anomalus, H. deserticola* and *H. paradoxus*. These were initially identified as hybrid species through early genetic studies that found they had a mixture of chloroplast DNA, allozyme and ribosomal DNA markers from both parental species (Rieseberg et al. 1990a; Rieseberg 1991). The hybrid species have strong ecological separation based on habitat choice; *Helianthus anomalus* occurs on sand dunes, *H. deserticola* on sandy soil of the desert floor and *H. paradoxus* in brackish salt marshes (Heiser et al. 1969). Comparative QTL mapping of hybrids and parental species suggested that this ecological separation was achieved by combinations of parental alleles that allowed for extreme (transgressive) phenotypes (Rieseberg et al. 2003). Subsequent experiments found that such transgressive phenotypes were favored when segregating hybrids of *H. annuus-H. petiolaris* were transplanted into the natural habitats of the three ancient hybrid species, with some synthetic hybrids rivaling the ancient hybrids in fitness (Lexer et al. 2003; Gross et al. 2004; Ludwig et al. 2004).

In addition to ecological isolation, the three hybrid species are partially reproductively isolated by hybrid pollen sterility caused mainly by chromosomal rearrangements (Lai et al. 2005). This was hypothesized to be due to the hybrid species having a combination of parental chromosomal rearrangements as well as unique changes (Rieseberg et al. 1995; Lai et al., 2005). Synthetic hybrids of *H. annuus-H. petiolaris* not only recovered fertility in a small number of generations, but also were more compatible with the putative hybrid species than were the parental species (Rieseberg et al. 1996; Rieseberg 2000, 2006). Microsatellite, chloroplast, and karyotypic analyses suggested all hybrid species originated between 63,000 and 208,000 generations before present and a single (*H. paradoxus*) or multiple (*H. anomalus* and *H. deserticola*) origins depending on the species (Schwarzbach and Rieseberg 2002; Welch and Rieseberg 2002; Gross et al. 2003). The strong link between admixture and reproductive isolation, in terms of ecology and karyotype, comparisons with experimental hybrid lineages, and partial replication across three species have made *Helianthus* a model for understanding homoploid hybrid speciation.

Although the *Helianthus* hybrid species were probed with cutting-edge molecular markers when initially identified, they have been relatively unexamined in the genomics age. Studies using Genotyping-By-Sequencing grouped *H. anomalus* and *H. deserticola* together in a phylogenetic tree but did not probe further (Baute et al. 2016). Moody and Rieseberg (2012) sequenced ten nuclear genes for multiple individuals of most annual sunflowers, including the hybrid species. They found extensive discordance between individual gene trees and notably that *H. anomalus* and *H. deserticola* grouped together and that *H. paradoxus* grouped with *H. annuus*, although the overall species tree did not agree with previous expectations. The most recent phylogeny of *Helianthus* did not include any hybrid species (Stephens et al., 2015). In each of these cases, modern approaches for identifying gene flow and hybrid ancestry were not performed. This is important because sunflower species are both young — *Helianthus* crown age is 3.6 mya — and generally have high effective population size (Strasburg and Rieseberg 2008). This means that incomplete lineage sorting (ILS) is likely high and discordant topologies that could be produced by hybridization may also be present without any interspecies gene flow. In addition, signals of hybridization can sometimes derive from unexpected sources (Owens et al. 2021).

Beyond the three well-characterized *Helianthus* hybrid species, numerous other cases of hybridization and introgression have been reported in the genus (Heiser et al. 1969; Rieseberg et al. 1991), leading Rieseberg (1991) to conclude that evolution in the group “must be considered reticulate, rather than exclusively dichotomous and branching.” While many of these early cases – which were based on relatively limited evidence – have been subsequently confirmed with genomic data (Kane et al. 2009; Owens et al. 2016; Mondon et al. 2018; Lee-Yaw et al. 2019; Zhang et al. 2019), others have not (Owens et al. 2021), and several remain to be tested. This highlights that the combination of modern genomic datasets and modern phylogenetic techniques allow for hypotheses about hybridization to be rigorously tested (Hibbins and Hahn 2022).

Here we use modern phylogenomic techniques to assess the extent of admixture during the evolution of the genus. To do this, we first use whole genome sequencing data for most annual sunflowers to infer the genome-wide phylogeny of this group. We then apply tests of introgression and gene flow across the annual clade, including the three hybrid species, *H. anomalus, H. deserticola* and *H. paradoxus*. This allows us to revisit criterion 2 of homoploid hybrid speciation for these three species, and to gain greater insight regarding their ancestry and their relationship to the broader patterns of admixture within the annual sunflower clade. Lastly, using a previously published phylogenomic data (Stephens et al., 2015), we extend our analyses across the entire genus.

## Results

### Data filtering

We generated whole genome sequence data for two *Helianthus paradoxus* samples. We called variants against the *H. annuus* HA412-HOv2 reference genome for these samples as well as previously sequenced data for two samples of eight other annual species, or subspecies, and four outgroups (supplementary table 1; Todesco et al. 2020). We retained 5,877,513 variable and 28,274,146 invariant sites after filtering for quality, mappability and call rate (supplementary figure 1). We divided the genome into a total of 11,797 10 kbp non-overlapping windows and retained 3,267 that had ≥ 2,000 called bases.

### Species and gene trees

The species topology for non-hybrid taxa from ASTRAL (fig. 1A,B) was mostly consistent with previous phylogenies that used fewer markers (Stephens et al. 2015; Baute et al. 2016; Zhang et al. 2019). Quadripartition support at nodes ranged from 38 to 84%, suggesting substantial variation in topology at gene trees. The maximum likelihood concatenated tree also found the same topology as ASTRAL and monophyly for all samples of the same species (fig. 1B).

**Figure 1:**
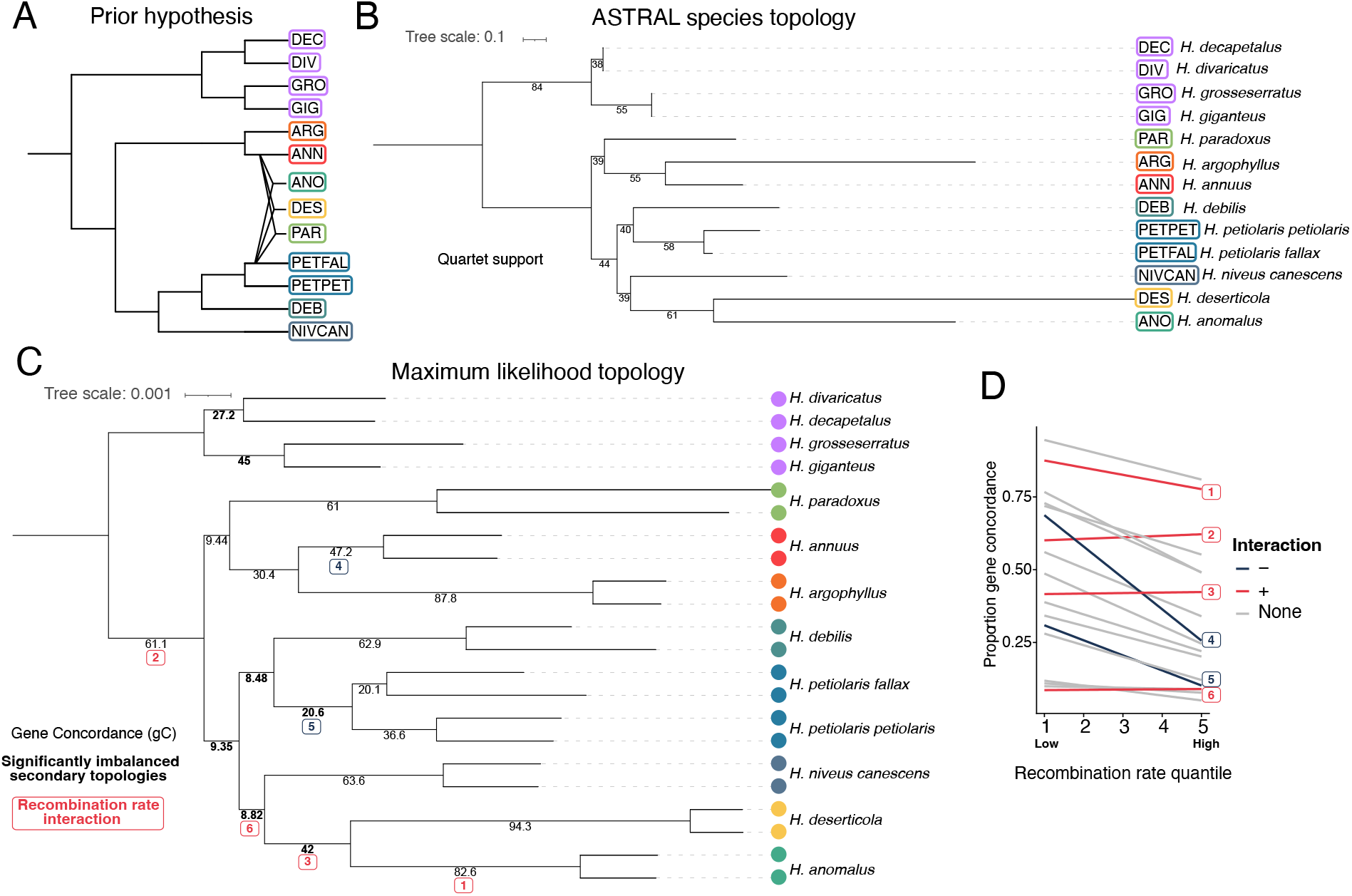
Phylogenetics of annual sunflowers. A) The prior hypothesis of annual species relationships (Rieseberg 1991; Stephens et al. 2015). B) Phylogenetic tree from combined 10 kbp gene trees used as input for ASTRAL species topology. Quartet support plotted on branches. C) Concatenated maximum likelihood tree using IQ-TREE2. Gene concordance plotted on nodes. Bolded values have significantly imbalanced secondary topologies. D) The relationship between gene concordance proportion and recombination quantile for each node with <90% overall gene concordance. Significant interactions with node ID are highlighted here and on panel C.

As outlined in the introduction, the three putative homoploid hybrid taxa, *H. paradoxus, H. deserticola* and *H. anomalus* are thought to have originated from hybridization between *H. annuus* and *H. petiolaris* (Rieseberg 1991). Since ASTRAL does not account for hybridization, we would expect the potential hybrid species to group near to the dominant parental species. We found that *H. paradoxus* grouped with *H. annuus* and its sister species *H. argophyllus*, whereas *H. anomalus* and *H. deserticola* grouped with *H. niveus canescens* (fig. 1B). Quadripartition support for these groupings is low (each 39%), as expected for hybrid lineages, although this was not unique to branches involving the previously identified hybrid species.

We further examined variation in gene tree topology by calculating the gene concordance factor for each branch on our concatenated tree using gene trees. We found relatively low support at branches separating species compared to branches separating samples within species (fig. 1C). When including all samples, each tree was unique. When we subsampled tips down to a single individual per species we found higher agreement; the most common tree was found 6 times out of 3,242 trees.

### Gene tree discordance

To explore why gene concordance values were low, we compared the proportion of trees matching the two alternate topologies for every quadripartition using IQ-TREE2’s gene concordance measure (Minh et al. 2020). Under purely incomplete lineage sorting, we expect that both alternate topologies should be found at equal proportions, while introgression will lead to imbalances. We found that at five nodes within the annual species, there was significant (X-squared, p<0.05) imbalance in alternate topologies (fig. 1C).

Selection against introgressed ancestry can lead to reduced introgression in regions of low recombination (Schumer et al. 2018; Martin et al. 2019). We explored this by using a binomial regression to quantify the relationship between support for the species topology and the recombination rate (as identified from the *H. annuus* genetic map) at each node with > 10% support for alternate topologies. In other words, are genomic regions with lower recombination more likely to show the species topology? We found a significant negative effect of recombination rate (β_1_ = -0.30, p = e^-14^) on support for the species topology (fig. 1C,D). Interestingly, we find a significant interaction between node and recombination rate at six nodes. At four nodes, most of which include the *H. anomalus/H. deserticola* clade, the relationship is less negative, or is positive (fig. 1D). At the node separating the *H. annuus* samples, the relationship with recombination is most strongly negative.

*Helianthus niveus canescens* was previously considered a variety of *H. petiolaris*, with which it intergrades for part of its range (Heiser et al. 1969). Our species topology places *H. niveus canescens* with *H. anomalus* and *H. deserticola* instead of *H. petiolaris*, but with relatively low support and overall less support at low recombination regions (fig. 1D). We counted the number of trees where *H. niveus canescens* grouped with *H. petiolaris petiolaris* and *H. petiolaris fallax* to the exclusion of *H. anomalus* and *H. deserticola*, as well as the reverse (supplementary figure 2). We found more support for *H. niveus canescens* together with *H. petiolaris*.

**Figure 2:**
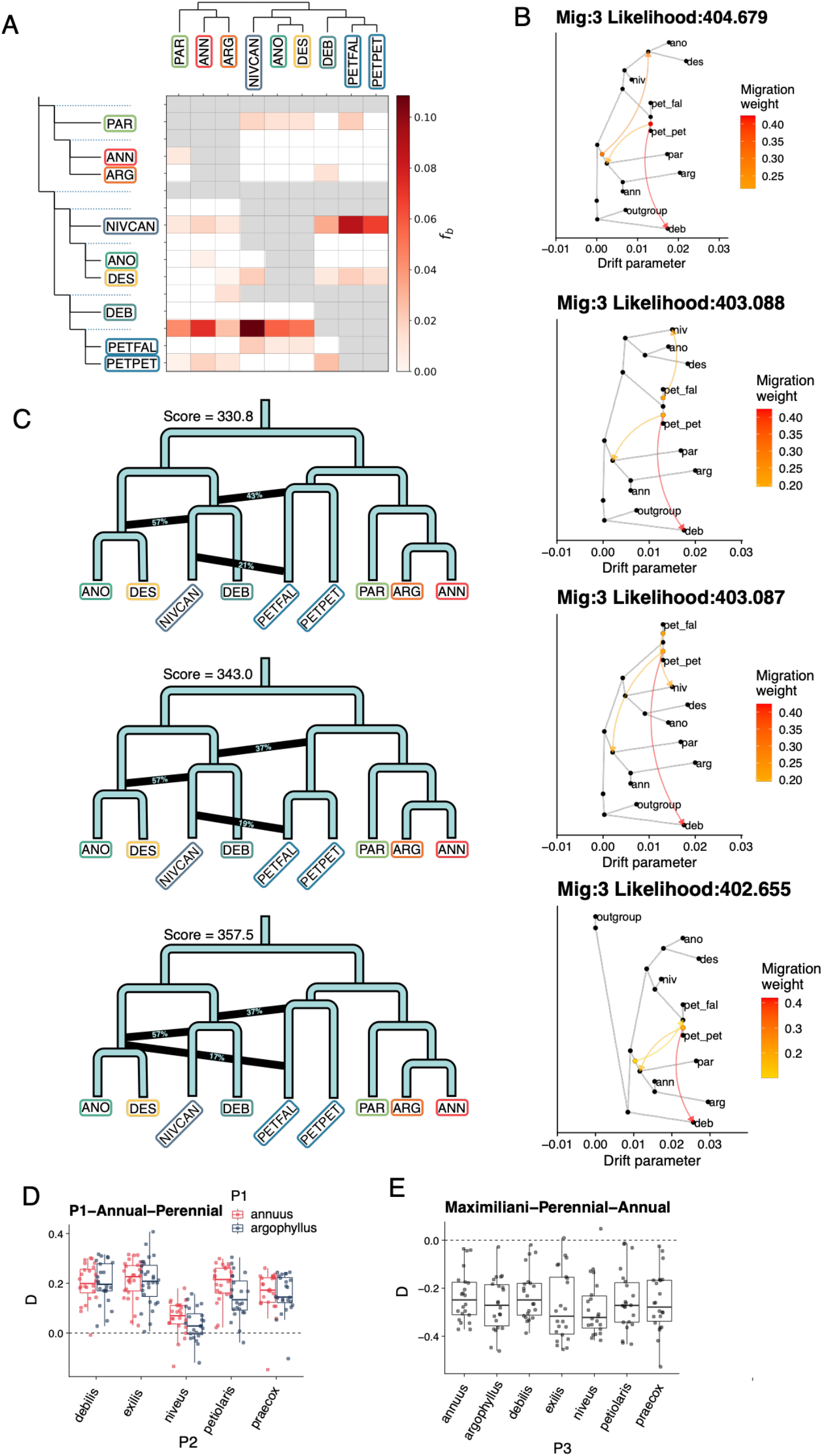
Introgression across sunflowers. A) F_branch_ statistic for annual species using the ASTRAL species topology. B) The four best supported TreeMix graph with three admixture edges. Higher likelihood score is better. C) The best supported admixture graph from ADMIXTOOLS2 with three admixture edges. Branch lengths are not to scale. Lower score is better. D) ABBA-BABA with the groupings [*H. annuus* OR *H. argophyllus*], other annual species, perennial species, *Phoebanthus*. Positive values indicate more ABBA counts. E) ABBA-BABA with the groupings *H. maximiliani*, other perennials, annual species, *Phoebanthus*.

### Admixture in the annual sunflowers

To assess the extent of hybridization during the evolution of the annual sunflowers, we used comprehensive approaches to test for admixture across all species. The f_branch_ analysis, which uses the f4-ratio test to identify introgression within a phylogenetic framework, found many significant signals (fig. 2A, supplementary table 2). The two strongest signals show that *H. petiolaris* is closer to all other species, when compared to *H. debilis*, and *H. niveus canescens* is closer to *H. petiolaris* and *H. debilis*, than *H. anomalus* and *H. deserticola* are. The f_branch_ patterns are dependent on the underlying phylogeny, so to explore this effect we repeated f_branch_ using the top nine most common species tree topologies (supplementary fig. 3). While no topology completely removes significant f_branch_ signal, this shows that introgression signals from f_branch_ can come from either close phylogenetic position or introgression. For example, in three of the top nine trees, *H. paradoxus* is placed within the *H. petiolaris* clade, but its f_branch_ signal shows it is much more similar to *H. annuus* and *H. argophyllus* than others in the *H. petiolaris* clade (supplementary fig. 3). The three hybrid species do not stand out in terms of the extent of admixture.

A different picture comes from the TreeMix analysis, which uses the covariance of population allele frequency to estimate a phylogeny and migration edges. We found that three was the optimal number of migration edges and explained 99.9% of variance. Above four edges, there was diminishing returns in likelihood gain and variance explained (supplementary figure 4). Admixture was found to be associated with the origins of the three putative hybrid species in the most likely TreeMix graphs at both three and four edges (supplementary figure 5). The origin of *H. paradoxus* appears to involve ancient hybridization between the ancestor of the *H. annuus* clade and *H. petiolaris*, although this admixture is inherited by the other sampled members of the *H. annuus* clade. Likewise, there is an admixture event between a basal lineage of the *H. annuus* clade and the ancestor of *H. deserticola* and *H. anomalus*. The third signal in common between the top three admixture graphs was between *H. debilis* and *H. petiolaris*. However, this likely is an artifact of misplacing *H. debilis* as basal in the tree and may instead suggest that it has ancestry from a lineage at the base of the annual sunflower clade. When we look other graphs that are close in likelihood to the top graphs, we frequently find admixture between *H. niveus canescens* and *H. petiolaris fallax*, although this results in the loss of the admixture event associated with *H. anomalus* and *H. deserticola*.

We also used ADMIXTOOLS2, which builds an admixture graph that explains f-statistics between populations. Admixture scenarios seen using ADMIXTOOLS2 have both similarities and differences with the TreeMix graphs. Out-of-sample scores decreased with increasing admixture edges from one to two, and from two to three, but not from three to four, suggesting that three admixture edges explain much of the f_2_ incongruence (supplementary figure 6). While bootstrapping was unable to distinguish between the top three graphs with three admixture nodes (p > 0.1), they each present a similar pattern that distinguishes them from the species tree (fig. 2C). The strongest signal is associated with origin of two of the putative hybrid species (*H. anomalus* and *H. deserticola*), but from admixture with the ancestor of *H. niveus canescens* and an unsampled or extinct basal lineage. *Helianthus petiolaris* is placed at the base of the *annuus* clade in the ADMIXTOOLS2 graph, implying that it shares ancestry with members of the *annuus* clade, including *H. paradoxus*. In addition, recent admixture is seen between *H. niveus canescens* and *H. petiolaris fallax* similar to that observed in the TreeMix analysis.

### Triplet tests of hybrid ancestry

While the admixture graphs (above) detected signals of hybridization associated with the origins of the three hybrid species, there were differences relative to previous hypotheses (Rieseberg 1991). Most notably, the origin of *H. anomalus* and *H. deserticola* appears to involve hybridization with the ancestor of *H. niveus canescens* rather *H. petiolaris fallax*. Also, there was uncertainty as to whether the signal of admixture associated with the origin of *H. paradoxus* was independent from other members of the *annuus clade*. Therefore, we explicitly tested hypotheses of hybrid ancestry for *H. anomalus, H. deserticola* and *H. paradoxus* against their putative parents. In the case of *H. paradoxus*, the testing is relative to *H. annuus* and *H. petiolaris fallax*, whereas for *H. anomalus* and *H. deserticola*, the putative parents are *H. annuus* and *H. niveus canescens*. For each test, we expect to see that the hybrid species is most closely related to the major parental species, and that tests of introgression show that the dominant signal of introgression involves the hybrid species and its minor parental species, rather than the parental species. We used four different approaches. Patterson’s D compares the amounts of shared derived alleles in a trio (Green et al. 2010; Malinsky et al. 2021). In this case, we are using Twisst to compare the counts of different gene tree topologies (Martin and Van Belleghem 2017). The branch length test compares the branch lengths in alternate topology gene trees. Lastly, QuIBL models ILS and introgression based on branch lengths. We note that the results of triplet tests should be interpreted with caution because of the long history of introgression between the putative parental clades (Yatabe et al. 2007; Strasburg and Rieseberg 2008; Kane et al. 2009). For the topology-based tests (Patterson’s D and Twisst), this means that the extent of introgression from the minor parent must be significantly greater than that between the parental lineages. For the distance-based tests (branch length and QuIBL), both the extent and the timing of introgression becomes critical, as the distance-based tests have little power for detecting ancient admixture.

For *H. paradoxus*, we consistently find *H. annuus* is the closest species, but inconsistent evidence of admixture with *H. petiolaris fallax* (fig. 3). Patterson’s D finds significant *H. paradoxus* – *H. petiolaris fallax* admixture. The results from Twisst are in the same direction, but are not significant. In contrast, the branch length test supports greater *H. annuus* – *H. petiolaris fallax* introgression and QuIBL supports neither (fig. 3). We find qualitatively similar results when using *H. petiolaris petiolaris* as a potential parental species. The branch length test is more sensitive to recent introgression, such as that ongoing between *H. annuus* and *H. petiolaris* (Yatabe et al. 2007), whereas D has greater power to detect ancient hybridization, perhaps accounting for the apparent discrepancy between the different tests. At the genomic window level, there is considerable variation in D with values ranging from -1 to 1 (supplementary figure 7).

**Figure 3:**
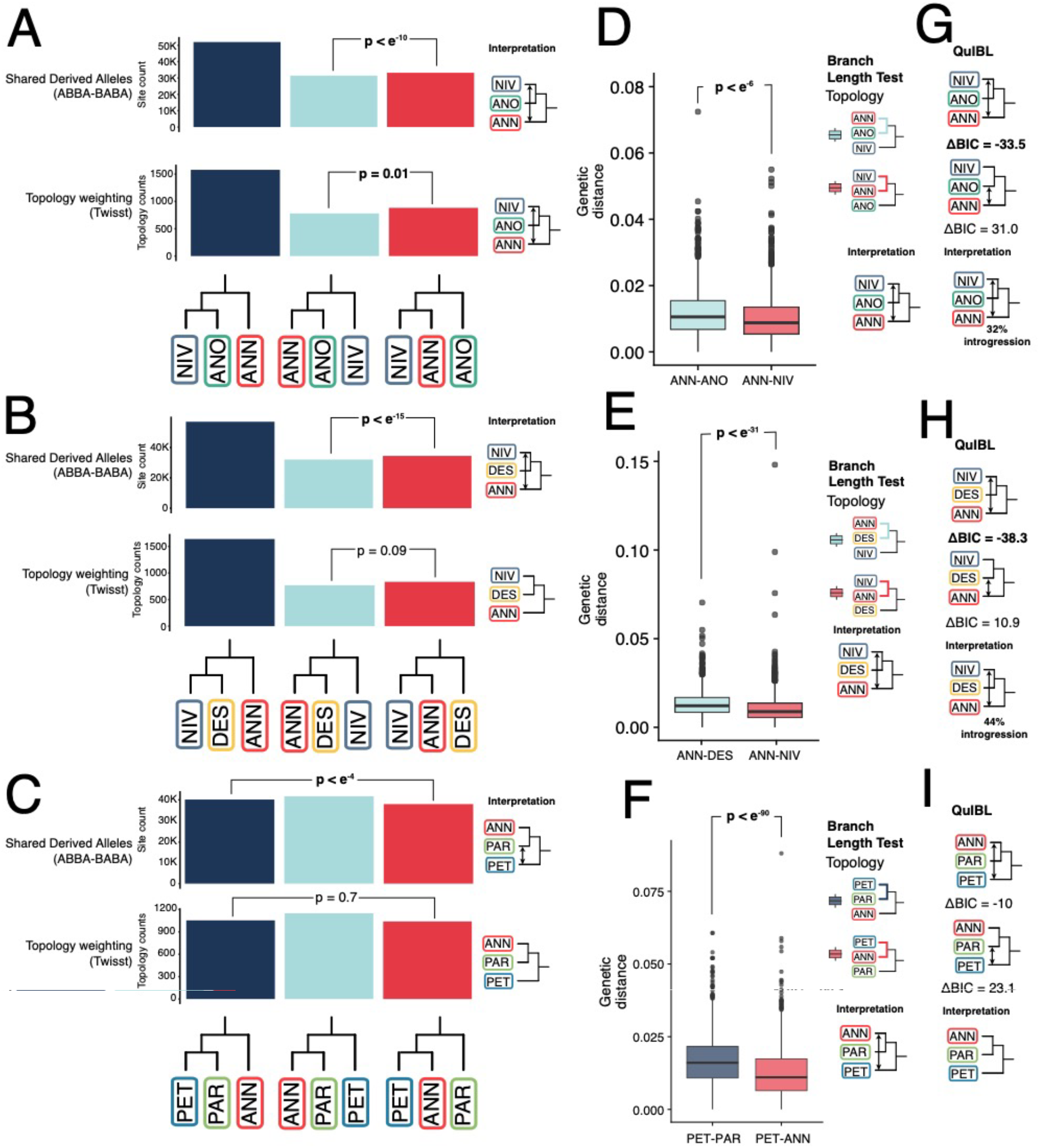
Triplet tests of previous hypotheses of hybrid ancestry. A-C) Counts of shared derived alleles and topological weighting for proposed hybrid species. P-values from block bootstrapping and chi-squared tests for ABBA-BABA and Twisst respectively. D-F) Branch Length Test of introgression showing genetic distance between taxa at gene trees with secondary topologies. G-I) QuIBL delta BIC values and interpretation. Lower values indicate more support for introgression.

For *H. anomalus* and *H. deserticola*, we see a consistent pattern for all four tests; they are more closely related to *H. niveus canescens* and there is more admixture between the potential parental species, than between the hybrid species and *H. annuus* (fig. 3). This could indicate that the hybrid species are ancient, and that the minor parent is at the base of the *H. annuus* clade (as suggested by TreeMix), and therefore has a relatively small contribution from contemporary *H. annuus*, or that the other parent of *H. anomalus* and *H. deserticola* is a now extinct basal lineage, as suggested in the ADMIXTOOLS2 graph.

### Admixture across the sunflower genus

To assess introgression across the rest of the genus, as well as to confirm our findings within the annual clade, we expanded our analysis of gene flow using a previously published sequence capture dataset spanning the genus. With this data, we used Dsuite to calculate f_branch_ and D for all trios (supplementary figure 8, supplementary table 3). We found evidence of introgression between perennial species, including *H. verticilliatus* – *H. giganteus, H. divaricatus* – *H. arizonensis* and *H. atrorubens* – *H. mollis*. This dataset also affirms the signal of introgression between *H. petiolaris* and both *H. niveus* and *H. annuus*. However, no introgression was found involving *H. debilis*. Surprisingly, we also find introgression-like signals between annual and perennial species. *Helianthus maximilliani* shares more derived alleles with annual species than other perennials (fig. 2E). Strangely, both *H. annuus* and *H. argophyllus* share fewer derived alleles with perennials than other annual species, which is consistent with widespread annual-perennial introgression excluding *H. annuus/H. argophyllus*, or introgression between the *H. annuus/H. argophyllus* clade and the outgroup *Phoebanthus* (fig. 2D).

## Discussion

For three-fourths of a century, *Helianthus* sunflowers have served as a model for the study of hybridization and its evolutionary role (Heiser 1947; Rieseberg 1991; Todesco et al. 2020). Early studies documented hybrid swarms between different sunflower species (Heiser 1947; Heiser 1949; Heiser 1951), demonstrated that chromosomal rearrangements contributed to hybrid sterility (Chandler et al. 1986; Lai et al. 2005), and speculated on possible ecotypes and species that might be the products of hybridization (Heiser 1949; Heiser 1958; Rieseberg 1991). Later molecular phylogenetic work and early low-resolution genomic studies supported some of these hypotheses (Rieseberg et al. 1990a; Yatabe et al. 2005), but not others (Rieseberg et al. 1988), and also suggested new hypotheses, including the putative origins of three homoploid hybrid species (Rieseberg 1991). Here, we have taken advantage of new high resolution genomic data and computational methods to further explore the role of hybridization in sunflower evolution. We pay particular attention to the three homoploid hybrid species, but we also examine other well-studied cases of hybridization in the *Helianthus* (and potential new cases) with the goal of reconciling the new data and analyses with previous work. Lastly, we explore the effects of variation in recombination rate on tree topology and introgression.

### Hybridization in sunflowers

The whole genome phylogeny for sunflower is consistent with that found by previous multi-locus studies (Stephens et al. 2015; Baute et al. 2016; Zhang et al. 2019). Leveraging this phylogeny, we employed several different methods to search for possible cases of admixture during the evolution of the genus. Remarkably, all species included in our study appear to be admixed, although the level of admixture depends in part on the particular analysis performed (fig. 2,4). Overall, the strongest signal of hybridization occurs between two subspecies of *H. petiolaris* and *H. annuus. Helianthus petiolaris* is sympatric with *H. annuus* and contemporary hybridization is well-documented (Heiser 1947; Rieseberg et al. 1998). Previous studies have documented “rampant” introgression between the species (Yatabe et al. 2005) and have suggested that interspecific gene flow was common in the past as well (Strasburg and Rieseberg 2008).

Another strong signal of admixture is found between *H. petiolaris* and *H. niveus canescens*, confirming a previous report (Zhang et al. 2019). *Helianthus petiolaris* intergrades with *H. niveus canescens* phenotypically, and Heiser et al. (1969) suggested that it might be due to hybridization. Although both types of phylogenetic analysis suggested *H. niveus canescens* grouped with *H. anomalus* and *H. deserticola* rather than *H. petiolaris*, there were more gene trees that placed *H. niveus canescens* with *H. petiolaris*. Considering the weight of evidence, we support that *H. niveus canescens* was involved with admixture of the *H. anomalus/deserticola* ancestor but was originally a member of the *H. petiolaris* clade. The beach sunflower, *H. debilis*, is sister to *H. petiolaris* in our tree, but f_branch_ shows that it is less similar to all other annual sunflower species (fig. 2A). This implies that it has ancestry from a lineage at the base of the annual sunflowers, as also suggested by TreeMix. Lastly, both the TreeMix and ADMIXTOOLS2 graphs suggest a shared hybrid origin for *H. anomalus* and *H. deserticola*, which we will discuss in more length below.

**Figure 4:**
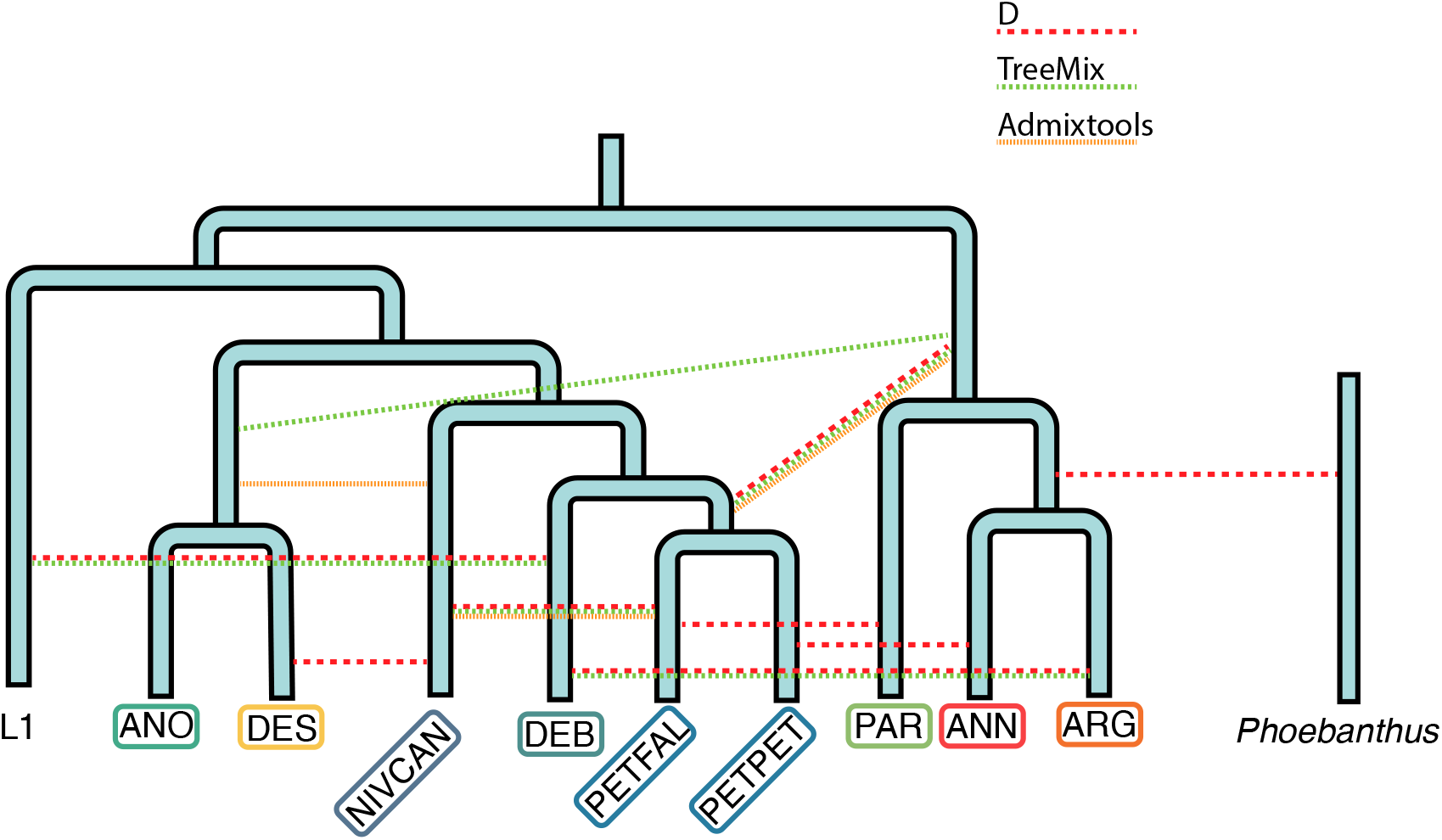
The proposed annual sunflower phylogeny with introgression events identified in the present study. L1 represents an unknown lineage seen through admixture analyses. Dotted lines indicate signatures of admixture seen in different analyses.

We used an alternate dataset generated by Stephens et al. (2015) to assess introgression across the entire *Helianthus* genus. We found evidence of hybridization between *H. verticilliatus* and *H. giganteus. Helianthus verticilliatus* was once thought to be a hybrid species but was later disproven by molecular data (Heiser et al. 1969; Ellis et al. 2006). Interestingly, *H. verticilliatus* was previously thought to be a subspecies of *H. giganteus*, and the two species overlap in range (Farwall 1916). Similarly, introgression between *H. mollis* and *H. atrorubens* is highly plausible considering they are known to hybridize (Beatley 1969). In contrast, the proposed introgression between *H. divaricatus* and *H. arizonensis* is difficult to explain because the species do not share any current range and there are no plausible proxy ancestry donors.

Surprisingly, there was identified introgression between the annual and perennial clades, despite strong post-pollination reproductive barriers between them. Most crosses between annual and perennial sunflower species fail to set seed, and the few successful hybrids that have been reported typically have low pollen viability (Heiser et al. 1969). Crosses involving the hexaploid perennials, represent an exception to this rule (Atlagić and Terzić 2006), but we found the strongest signal of introgression with a diploid perennial *H. maximiliani*, which is incompatible with members of the annual clade (Henn et al. 1998). Thus, we likely are seeing the outcome of ancient hybridization, which occurred before reproductive isolation was complete. However, the most unexpected pattern is a signal of ancestral allele sharing between *H. annuus/argophyllus* and the outgroup, *Pheobanthus* (fig. 2E). These two lineages diverged 2.5-5.4 mya, and are therefore expected to be completely reproductively isolated (Owens and Rieseberg 2014; Mason 2018), again suggesting that the signal of admixture is from ancient hybridization with a now extinct species. Nonetheless, we recommend experimental crosses be undertaken between *Pheobanthus* and members of the annual clade to see if hybrids can be made. It is also possible that this signal represents a change in evolutionary rate in *H. annuus* and *H. argophyllus*. This has been proposed to explain similar patterns of introgression signals in *Papilio* butterflies (Xiong et al. 2022). More broadly, our results suggest that admixture has been common in *Helianthus* throughout its evolutionary history, and that it has had a profound effect on phylogenomic relationships in the genus (fig. 4). Thus, *Helianthus* can be viewed as a syngameon, in which gene flow connects otherwise distinct species (Grant 1981). Our findings also address earlier concerns that hybridization in *Helianthus* might be a recent consequence of anthropogenic disturbance and therefore of little significance to its long-term evolution (Schemske 2000). Whether these ancient hybridization events triggered diversification, as suggested in other groups (e.g., Meier et al. 2017), remains unclear.

### Ancestry of putative homoploid hybrid species

The TreeMix and ADMIXTOOLS2 graphs indicate that the three hybrid species are admixed, thereby fulfilling the second criterion for homoploid hybrid speciation. However, there are differences relative to the original scenario put forward by Rieseberg (1991). In particular, the hybridization events associated with the origins of the hybrid species appear to have occurred further back in time than previously believed. For *H. paradoxus*, a potential hybrid speciation event is older than previously anticipated (Welch and Rieseberg 2002) since individual gene trees place *H. paradoxus* most often at the base of the annuus clade, therefore at least older than 1 mya. For *H. anomalus* and *H. deserticola*, our phylogenomic results suggest they are sister species, and share an admixture close in time to their origin. This would also place a potential hybrid speciation event further in the past than previous work suggested (Schwarzbach and Rieseberg 2002; Gross et al. 2003; see supplementary discussion for reconciliation with previous microsatellite dating of the hybrid species origin).

Another important difference is that *H. anomalus* and *H. deserticola* are sister species and their major parent appears to be *H. niveus canescens* rather than *H. petiolaris fallax* as originally proposed (Rieseberg 1991). While this was unexpected, *H. niveus canescens* intergrades phenotypically with *H. petiolaris fallax* and the two taxa share chloroplast and nuclear ribosomal DNA haplotypes, which were the markers employed to hypothesize a hybrid origin in the first place (Heiser et al. 1969; Beckstrom-Sternberg et al. 1991). As a consequence, some taxonomic treatments have considered *H. niveus canescens* to be a variety of *H. petiolaris* (Blake 1942; Schilling 2020). The present study indicates that *H. niveus canescens* is genetically distinct from *H. petiolaris fallax*, although there is considerable admixture between them, which might account for their phenotypic resemblance (Zhang et al. 2019). Fortunately, recognition that the ancestor of *H. niveus canescens* was a parent of *H. anomalus* and *H. deserticola* has little impact on the interpretation of previous work. The population of *H. petiolaris fallax* that was employed for most of the experimental studies of hybrid speciation came from the zone of intergradation with H. *niveus canescens* (Rieseberg et al. 2003; Lexer et al. 2003; Gross et al. 2004; Ludwig et al. 2004). The taxa are indistinguishable morphologically and ecologically in this area, and may recapitulate the ancestor of *H. niveus canescens*.

Despite the power of the phylogenomic analyses, some uncertainties remain (see supplementary discussion about the challenges for interpreting admixture signals in sunflower). For *H. paradoxus*, it is unclear whether the signal of admixture associated with its origin is independent from other members of the *annuus clade*. The triplet tests of introgression provide indirect support for independence. For D, there is a significant enrichment for *H. petiolaris* – *H. paradoxus* sharing as predicted under the hybrid origin scenario; gene tree topology counts show the same pattern but are not significant. However, the branch length test, which has greater power to detect recent introgression, indicates more introgression between *H. annuus* – *H. petiolaris*. These results may be reconciled by the long history of introgression between the *annuus* and *petiolaris* clades (Strasburg and Rieseberg 2008), with the branch length test successfully detecting recent introgression between *H. annuus* and *H. petiolaris*, and D offering evidence of more ancient hybridization associated with the evolution of *H. paradox*. Likewise. *H. paradoxus* shares more derived alleles with *H. anomalus* and *H. deserticola* than do other members of the *annuus* clade, offering further indirect support that *H. paradoxus* is the product of an independent hybridization event (supplementary table 2).

There is also uncertainty regarding the parentage of *H. anomalus* and *H. deserticola*. While the role of *H. niveus canescens* as one parent is strongly supported, the other hybrid parent is less clear. Is it the *H. annuus* lineage? The most likely TreeMix graph suggests the ancestor of the *H. annuus* clade is the other parent, but this pattern is not strongly supported by admixtools, the triplet testing or D. The hybrid species share fewer derived alleles with *H. annuus* and *H. argophyllus* when compared to *H. debilis, H. niveus canescens* or *H. petiolaris* (supplementary table 2), but they share more derived alleles with *H. paradoxus* when compared to *H. debilis*.

This may be indicative of more recent introgression between the *H. annuus* lineage and members of the *petiolaris* clade sampled in this study, all of which overlap in geographic distribution and hybridize with *H. annuus*. Alternatively, *H. anomalus* and *H. deserticola* are a combination of *H. niveus canescens* and an unsampled or extinct basal lineage, as is suggested in the admixtools tree. Future comparisons with an allopatric control from the *H. petiolaris* clade (e.g., *H. niveus niveus*) will be needed to distinguish between these hypotheses.

One perhaps surprising aspect of our results is that the amount of admixture in “hybrid species” is not exceptional compared to non-hybrid species. This highlights that the amount of hybrid ancestry is not critical for hybrid speciation, only that admixture was involved in reproductive isolation (Rieseberg 1997; Schumer et al. 2014), which we do not directly address in this study. Nonetheless, our results have implications for the interpretation of previous experimental studies that have attempted to make this link (e.g., Rieseberg et al. 1995, 1996, 2003; Lexer et al. 2003; Gross et al. 2004; Ludwig et al. 2004). For example, evidence that the homoploid hybrid species are more ancient than previously believed makes it more difficult to determine which ecological, phenotypic and genomic changes are associated with hybridization and which changes arose as a consequence of divergence after hybrid speciation (Ungerer et al. 2006). We suspect that the clusters of parental markers found to be linked to QTLs in segregating hybrids, and which were also found in the hybrid species genomes (Rieseberg et al. 2003), likely correspond to inversions. Such inversions are frequent in *Helianthus* (Ostevik et al. 2020) and often appear to be ancient, pre-dating the species they are found in (Todesco et al. 2020). Future studies attempting to connect reproductive barriers with admixture should be conducted within a phylogenetic framework to better assess when key differences arose.

Previous work attempted to dissect parental ancestry across the putative hybrid genomes, finding strings of linked parental markers suggestive of rapid establishment of the hybrid genomes (Ungerer et al. 1998; Buerkle and Rieseberg 2008). However, incomplete lineage sorting, multiple ancient admixture events, structural variation, and the auto-correlational effects of recombination rate variation may all affect ancestry assignments across the genome. Future attempts at identifying and interpreting ancestry tracks will have to contend with this complexity.

Are there other examples of homoploid hybrid speciation in *Helianthus*? Probably – although the relevant data come from earlier studies rather than the present analysis (see supplementary discussion about outcomes of hybridization in the annual sunflower clade). Independently derived dune ecotypes of the prairie sunflower are isolated from non-dune populations by multiple reproductive barriers (Heiser 1958; Ostevik et al. 2016). Recent population genomic studies indicate that most of the traits underlying reproductive isolation map to ancient chromosomal inversions that appear to have originated via introgression from a basal lineage in the annual clade (or even earlier) (Huang et al. 2020; Todesco et al. 2020). In an example involving more recent hybridization, a 77-day difference in flowering between coastal and barrier island ecotypes of *H. argophyllus* was found to result from a 30 Mb introgression from *H. annuus* containing a functional copy of *HaFT1* (Todesco et al. 2020). The dune ecotypes arguably represent new species, and one of them is described as *H. neglectus* in most taxonomic treatments (Heiser et al. 1969; Schilling 2020). Ecotypic differentiation in *H. argophyllus* represents an earlier stage of the speciation process, but it illustrates how introgression of a single gene can cause significant reproductive isolation. It is noteworthy that all three potential new cases of homoploid hybrid speciation involve the colonization of ecologically divergent habitats that provide ecogeographic isolation similar to that seen in the ancient hybrids.

### Recombination rate and introgression

Local recombination rate is a critical population genetics parameter that can affect ILS, gene flow and genetic diversity (Ortíz-Barrientos et al. 2002; Nachman and Payseur 2012; Haenel et al 2018). For example, introgressed loci are more likely to persist in regions of high recombination because they more quickly unlink from deleterious neighbouring loci, such as genetic incompatibilities (Brandvain et al. 2012; Schumer et al. 2018). In our dataset, higher recombination regions are less likely to support the inferred species topology. This is likely a combination of reduced introgression in low recombination regions, as well as lower Ne, which reduces ILS (Martin et al. 2019; Li et al. 2019; Pease and Hahn 2013). The effect is strongest for the node separating *H. annuus* samples, where the recombination rate estimates are most accurate. Sunflowers have highly labile chromosome structure and there are numerous large-scale rearrangements between species in this tree, but this pattern is retained even between our outgroup perennial sunflower species, suggesting that recombination rate is relatively conserved even if genome structure changes (Ostevik et al. 2020). At several nodes we find that the relationship is reversed, which may be because at those nodes we have the introgressed topology, rather than the true species topology. Exploring the relationship between recombination rate and tree topology is useful for identifying an accurate species topology.

### Conclusions

Despite vast improvements in the quantity of genetic data available to current researchers, disentangling introgression patterns is often challenging (Hibbins and Hahn 2022). When multiple introgression events have occurred, this can lead to false positives or false negatives, which is especially true when the true donors are not sampled or, in some cases, no longer extant. Broad sampling of species can help reduce these issues and broad sampling of populations within species can identify when introgression is geographically localized. Thus, when evaluating the plausibility of introgression signals detected using genomic data, it is important to consider factors such as reproductive compatibility and the likelihood of geographic overlap over the course of evolutionary divergence. That being said, the reconstruction of historical ranges is an inexact science, and information on extinct congeners is completely unknown for most wild genera.

Phylogenomic analysis below the genus level often finds evidence of introgression, especially in plants (Dagilis et al., 2021). This suggests that a significant proportion of plant species have some admixed ancestry. Despite this, homoploid hybrid species are rare because most studies fail to link admixture with the evolution of reproductive barriers (Schumer et al., 2014). Although attempts have been made to identify admixture-derived reproductive barriers through purely bioinformatic analyses (e.g. Sun et al. 2020; Wang et al. 2022), links between phenotypes, reproductive isolation and introgressed ancestry are required to confirm the homoploid hybrid speciation. A notable exception to this is work by Wang et al. (2021), which identified an admixed species in *Ostryopsis*, quantified reproductive barriers and transgenically tested candidate genes acquired through admixture. Widespread introgression across sunflower species raises the possibility that admixture played a role in other speciation events. In particular, chromosome rearrangements are both common and are known to cause reproductive isolation in sunflowers (Ostevik et al. 2020; Lai et al. 2005), so future work should leverage chromosome resolved genomes and QTL mapping to identify how whether the introgressed rearrangements cause speciation.

## Methods

### Data generation

We generated whole genome resequencing data for two *H. paradoxus* samples. DNA was extracted from individual seedlings or dried leaf tissue using a modified CTAB extraction protocol (Murray et al. 1980; Zeng et al. 2002), and an indexed Illumina sequencing library was created, which included a Duplex Specific Nuclease (DSN) repeat depletion step (Todesco et al. 2020). This data was added to previously published whole genome resequencing data to create a dataset of nine species, or subspecies, of annual sunflowers (Hubner et al. 2019; Todesco et al. 2020; Owens et al. 2021). For each, we randomly selected two samples to be used in the analysis. We only included two per species because we were interested in understanding between species relationships, rather than within species diversity. Additionally, using fewer samples substantially reduced the computational bottleneck in both variant calling and subsequent phylogenetic analyses. To act as outgroups, we also included one sample each for four diploid perennial sunflower species (See supplementary table 1).

For each sample, Illumina sequence data was aligned to the *Helianthus annuus* HA412-HOv2 genome using NextGenMap (v0.5.3) (Sedlazeck et al. 2013), PCR duplicates were marked with samtools v0.1.19 (Danecek et al. 2021), and variants were called using GATK (v4.1.4.1) (Mckenna et al. 2010). We specifically included invariant sites in GATK genotypeGVCFs to facilitate downstream analyses. The sunflower genome contains a large proportion of transposable element repeats that hamper read alignment. To avoid those regions, we used GenMap (-K 50 -E 2) (v1.3) to calculate k-mer uniqueness or mappability for all positions in the genome (Pockrandt et al. 2020). We then removed all contiguous regions with mappability < 1 that are greater than 100 bp. This removed 2.43 GBp out of 3.23 Gbp, but should highly enrich retained regions for single copy sequences.

For our dataset, we filtered variant and invariant sites separately. For variant sites, we removed all indels and visualized the distribution of quality metrics of SNPs using bcftools (v1.14) to extract values and ggplot (v3.3.5) in R (v4.1.2) to plot (Whickham 2011; R core team 2021). We visually selected thresholds to remove outliers based on the distribution (bcftools view -e ‘INFO/DP > 1000’ -e ‘INFO/FS > 50’ -e ‘INFO/QD < 2’ -e ‘INFO/SOR > 4’ -e ‘INFO/MQ < 30’) (supplementary figure 1). Variant and invariant sites were combined, and then filtered to require ≥ 4 reads per genotype and ≥ 80% of samples genotyped using VCFtools (v0.1.16) (Danecek et al. 2011).

Code used to conduct analyses and make figures is deposited at https://github.com/owensgl/helianthus_hybrid_species_2021.

### Gene and species tree creation

We took two approaches to explore genome-wide gene trees. In the first, we equalized information across windows by requiring each genomic window have 10,000 called bp, leading to variable physical size. In the second approach, we equalized physical size but not information by dividing the entire genome into non-overlapping 10,000 bp windows and retained windows with ≥ 2,000 called bases. In both cases, heterozygous sites were retained and coded using IUPAC coding.

For each of our genomic windows, we calculated a maximum likelihood phylogeny using IQ-Tree (v2.0.6) (Minh et al. 2020a) with *H. giganteus*, a perennial species, as the outgroup. Additionally, a single concatenated species tree using all retained bases was created using maximum likelihood and model selection in IQ-TREE. We used a coalescent approach in ASTRAL (v5.7.3) to estimate the species tree using both sets of gene trees separately (Zhang et al. 2018).

We explored the diversity of gene tree topologies by using the tool findCommonTrees.py (Edelman et al. 2020) to identify and count the occurrence of each topology. We found that every tree was unique, so we subsampled trees down to a single sample per species and a single perennial outgroup species to reduce the possible tree space. We repeated the counts of trees and visualized the most common gene trees.

Using the concatenated phylogeny as a backbone, we measured gene concordance using IQ-Tree2 (Minh et al. 2020b). This measures the percent of gene trees that support the topology and the two possible alternate topologies for each branch quartet. We matched the *H. annuus* recombination rate for each genomic window and calculated the percent support for each topology for each recombination quintile. To determine if recombination rate affected support for the species topology, we used a binomial regression with the formula:

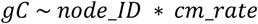

### Broad tests of introgression

We used several methods that integrate signals of introgression between all samples. Using Dsuite, we calculated the f_branch_ statistic for the species phylogeny (Malinsky et al. 2021). This statistic incorporates all valid f_4_ ratios (similar to Patterson’s D) as well as the phylogeny to identify branches that share allelic imbalance. This can help identify introgression events that predate speciation times and therefore have a signal in multiple species.

We also used TreeMix to identify introgression events (Pickrell and Pritchard 2012). This program builds a graph model of population splits based on the covariance of allele frequencies. It allows for migration edges to be added to the graph that explain remaining covariance not accounted for by the initial graph topology. Since TreeMix relies on covariance, we pruned our dataset for SNPs in LD by only including sites with > 0.8 r^2^ in 100,000 bp using ‘bcftools prune’ (Danececk et al. 2021), and did not include sample size correction. We ran TreeMix with 0 to 10 migration edges, with 99 replicates using different starting seeds, and selected the optimal number of migration edges using the Evanno method in OptM (Fitak 2021). We visualized the replicates with the likelihood values within 10 of the highest likelihood for each number of migration edges (i.e. the models that were not the best, but were close).

Lastly, we built an admixture graph for the phylogeny using ADMIXTOOLS 2 (Patterson et al. 2012). We first converted our VCF file to eigenvector format using a custom Perl script, and then calculated all f2 statistics using the extract_f2 and f2_from_precomp functions. We used a 5 Mbp window for block bootstrapping. Admixture graphs were found using the find_graphs command with stop_gen=200, stop_gen2=20 and perennials set as the consistent outgroup. This method finds graphs that fit the observed f-statistics in the data. The graph search can get stuck at local optima, so we repeated each search 100 times for each number of admixture events, from one to four, and retained the best fitting graph in each iteration. From this, we picked the top three scoring graph for each number of admixture events. We then used bootstrap-resampling of the out-of-sample scores for each graph using the function qpgraph_resample_multi to ask whether adding additional nodes was significantly better supported, by comparing the best graph with n admixture events with the best graph with n+1 admixture events, as well as comparing graphs.

### Trio based tests of introgression

Given the high levels of discordant gene trees, and conflicting signals from broad tests of introgression, we explored trio-based tests of introgression using Dsuite to calculate Patterson’s D (Green et al. 2010; Malinsky et al. 2021). This test looks at quartets with the relationship (((A,B),C),O) and asks if there is greater derived allele sharing between A and C or B and C. Under ILS, both should share equal counts of derived alleles, but introgression will lead to imbalance. Significance was tested using a block bootstrap algorithm. We used the perennial species as an outgroup and tested all trio sets consistent with the species phylogeny.

For the putative hybrid species and their parents, we used three additional approaches, the branch length test (BLT), QuIBL and Twisst. For these tests, we used genomic windows with equal physical size, as described above. The BLT test compares the tip-to-tip distance between inferred sister clades in the minor topologies using a Mann-Whitney test (Suvorov et al. 2022). Under ILS, both minor topologies should have equal distance, but if introgression is occurring then the topology it produces should have reduced distance due to its more recent coalescence. These tests were done in R using TreeTools (Smith 2019), using the subsampled trees with only a single sample per species. QuIBL examines trios in gene trees and asks whether the branch length distribution fits better to a model that includes both ILS and later introgression, or purely ILS (Edelman et al. 2020). As before, we used our perennial samples as outgroups. We considered models with Bayesian information criterion score difference of ≥ 30 to be evidence of introgression. We used Twisst to calculate the topological weighting for each possible tree using the equal size gene trees (Martin and Van Belleghem 2017). This takes into account phylogenetic position variation amongst individuals within a species and calculates a weight for how common a particular topology is at each gene tree. We normalized weights such that each genomic window had a total weight of one and counted the total weight for each topology. Similar to the Discordant Count Test from Suvorov et al. (2020), we compared the counts of the two minor topologies using a chi-squared test, under a null hypothesis that ILS should produce equal counts of both minor topologies. We also calculated D across the genome using Dsuite, with a window size of 50 SNPs and a step size of 25 for each possible candidate trio.

### Quantifying introgression across the genus

Given the amount of gene flow identified within annual sunflowers, we expanded our analysis of gene flow by using a previously published sequence capture set encompassing a majority of diploid species in the genus (Stephens et al. 2015). This dataset included 170 aligned sequences totalling 106,862 bp and 11,407 parsimony informative sites. Since these markers were anonymous and genome positions are required for block bootstrapping, we aligned them to the *H. annuus* HA412-HOv2 reference genome using BLAST and selected the location hit with the highest e-value. Sequence capture gene alignments include numerous indels, so we were not able to combine this dataset with our previous WGS dataset. As before, we used Dsuite to calculate D, f_4_ and f_branch_. For the species topology, we used the MP-EST tree from Stephens et al. (2015), with polytomies randomly resolved.

## Supporting information

Supplementary discussion and figures

## Acknowledgements

We would like to thank Sarah Yakimowski and Robert Sivinski for providing tissue used for DNA extractions. This work was funded by a Banting Fellowship to GLO, as well as a Discovery Grant from the Natural Sciences and Engineering Research Council of Canada to LHR.

