## Supplementary discussion and figures for "Re-evaluating homoploid reticulate evolution in the annual sunflowers"

Gregory L. Owens[1*]

Kaichi Huang[2]

Marco Todesco[2]

Loren H. Rieseberg[2]

[1]: Department of Biology, University of Victoria, BC, Canada

[2]: Department of Botany and Beaty Biodiversity Center, University of British Columbia, BC, Canada

### *Supplementary Material*

#### *Supplementary Discussion.*

##### *Age of hybrid species*

Our phylogenomic analyses suggest an earlier origin for the three hybrid species than did previous studies (Welch and Rieseberg 2002; Schwarzbach and Rieseberg 2002; Gross et al. 2003), which used divergence at microsatellite loci to tentatively place their origins at between 210,000 and 63,000 generations before present. However, earlier authors were appropriately cautious about these estimates. For example, Welch and Rieseberg (2002) cautioned that their estimate should "be viewed with considerable skepticism" because of uncertainty about microsatellite mutation rates in sunflower. Also, keep in mind that sunflowers, while annual, have a seed bank, which can cause elevated generation times. For example, if we were to assume generation times of 3-4 years, similar to that recently reported for *Arabidopsis thaliana* (Lundemo et al. 2009), this would push the origin of the three species back in time, more consistent with the present study. Desert annuals (like *H. anomalus* and *H. deserticola*) have notably strong seed dormancy mechanisms, likely suggestive of even longer generations. The previous studies (Welch and Rieseberg 2002; Schwarzbach and Rieseberg 2002; Gross et al. 2003)) concluded that the origin of the hybrid species pre-dated the arrival of humans in North America, as also suggested by our results.

#### *Challenges for interpreting admixture signals in sunflower*

Part of the challenge of determining the speciation and introgression history of sunflowers is the high percentage of anomalous gene trees (Moody and Rieseberg 2012). For almost all nodes separating species, the species topology was not the majority gene tree topology (fig. 1C). One reason for this is that some sunflower species have high effective population sizes (Strasburg et al. 2011b) and the genus speciated relatively rapidly in the last four million years (Mason 2018). Consequently, detecting specific introgressed loci is challenging because gene trees matching introgression scenarios are common, even when genome-wide signals of introgression are absent (Zheng and Janke 2018). Gene concordance at nodes separating individuals within a species represent the proportion of gene trees where species are monophyletic. These values varied for individual species from 94% in *H. deserticola* to 20% in *H. petiolaris fallax*. Effective population size and recent introgression will affect gene tree monophyly, but are partially confounded by differences in species sampling. Samples were randomly selected from previously sequenced sets, and some species had much greater sampling across their range. For example, both *H. debilis* samples were from Texas, while the entire species spans a much larger region, and therefore we may be overestimating monophyly for the species. Lastly, detected admixture may come from extant or extinct species that were not included in the present study, potentially leading to faulty conclusions.

#### *Outcomes of hybridization*

Hybridization can have several different outcomes, including the evolution of stable hybrid zones, introgression, reinforcement, ecotype formation and speciation, and the merger of previously independent populations (Abbott et al. 2013; Todesco et al. 2016). While we do not yet know the frequency of these different outcomes, both theoretical and empirical studies suggest that the formation of stable hybrid zones and introgression are more frequent outcomes of hybridization than hybrid speciation or extinction (Barton and Hewitt 1985; Barton 2001; Buerkle et al. 2003; Abbott et al. 2013; Todesco et al. 2016; Owens and Samuk 2020). Homoploid hybrid speciation only becomes likely when there is an open habitat available for the hybrids that affords some level of spatial and ecological separation for the new hybrid lineage (Buerkle et al. 2003). This is what we see in *Helianthus*; hybridization between widespread taxa with broadly overlapping distributions such as *H. annuus – H. petiolaris* and *H. petiolaris – H. niveus canescens* has resulted in high levels of introgression and phenotypic intergradation. In addition, introgression has been reported between in zones of contact between *H. annuus* and local endemics such as *H. bolanderi* in California (Owens et al. 2016) and *H. argophyllus* in Texas (Owens et al. 2021). Only when such hybrids are able to colonize a divergent habitat that is spatially isolated from congeners (such as in the three homoploid hybrid species), does admixture appear to trigger speciation. Note that all other annual sunflowers included in this study overlap geographically and currently hybridize with other congeners, further highlighting the uniqueness of the homoploid hybrid species.

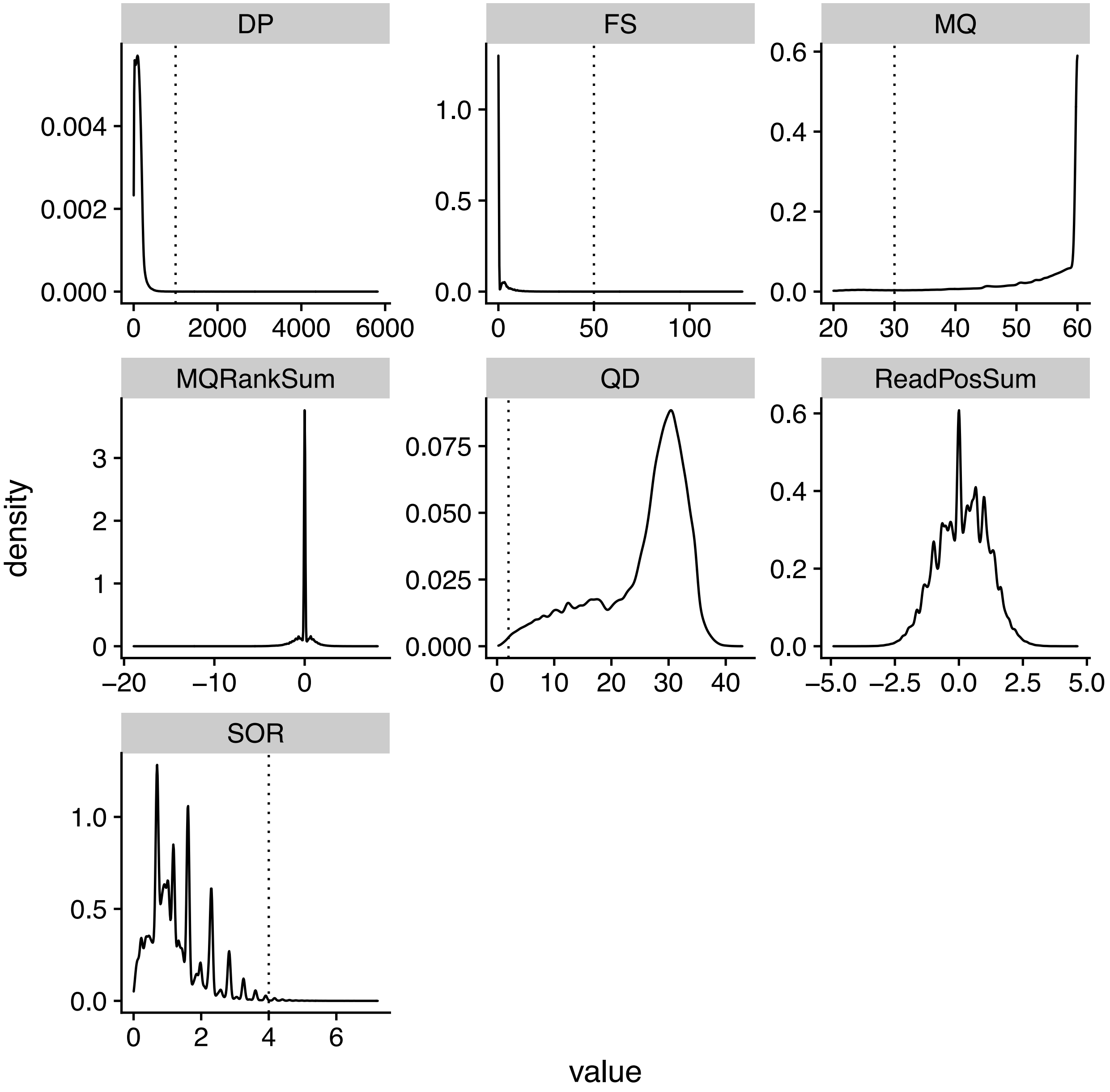


**Supplementary Figure 1: The distribution of quality scores for called variants.** Cutoff values used for filtering are shown with the dotted line.

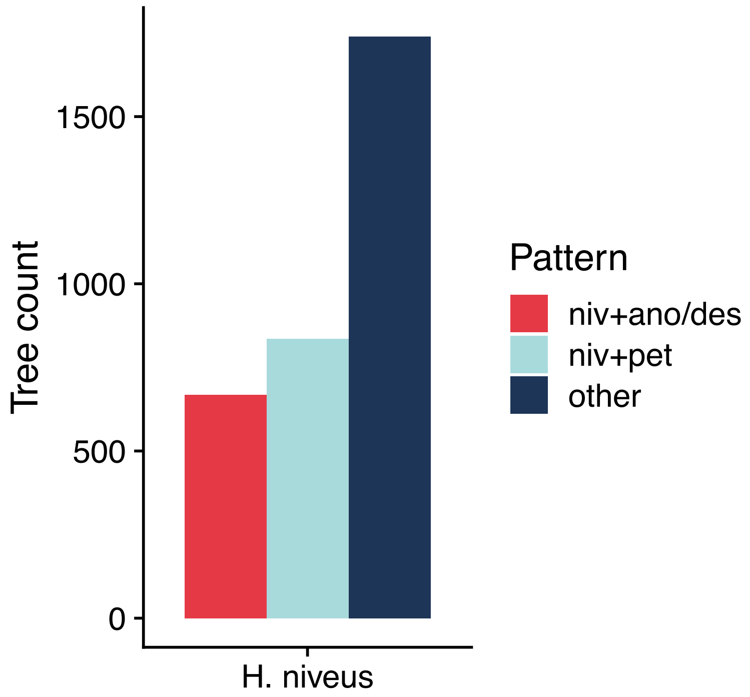


**Supplementary Figure 2: Counts of topologies for gene trees generated from 10 kbp windows.**

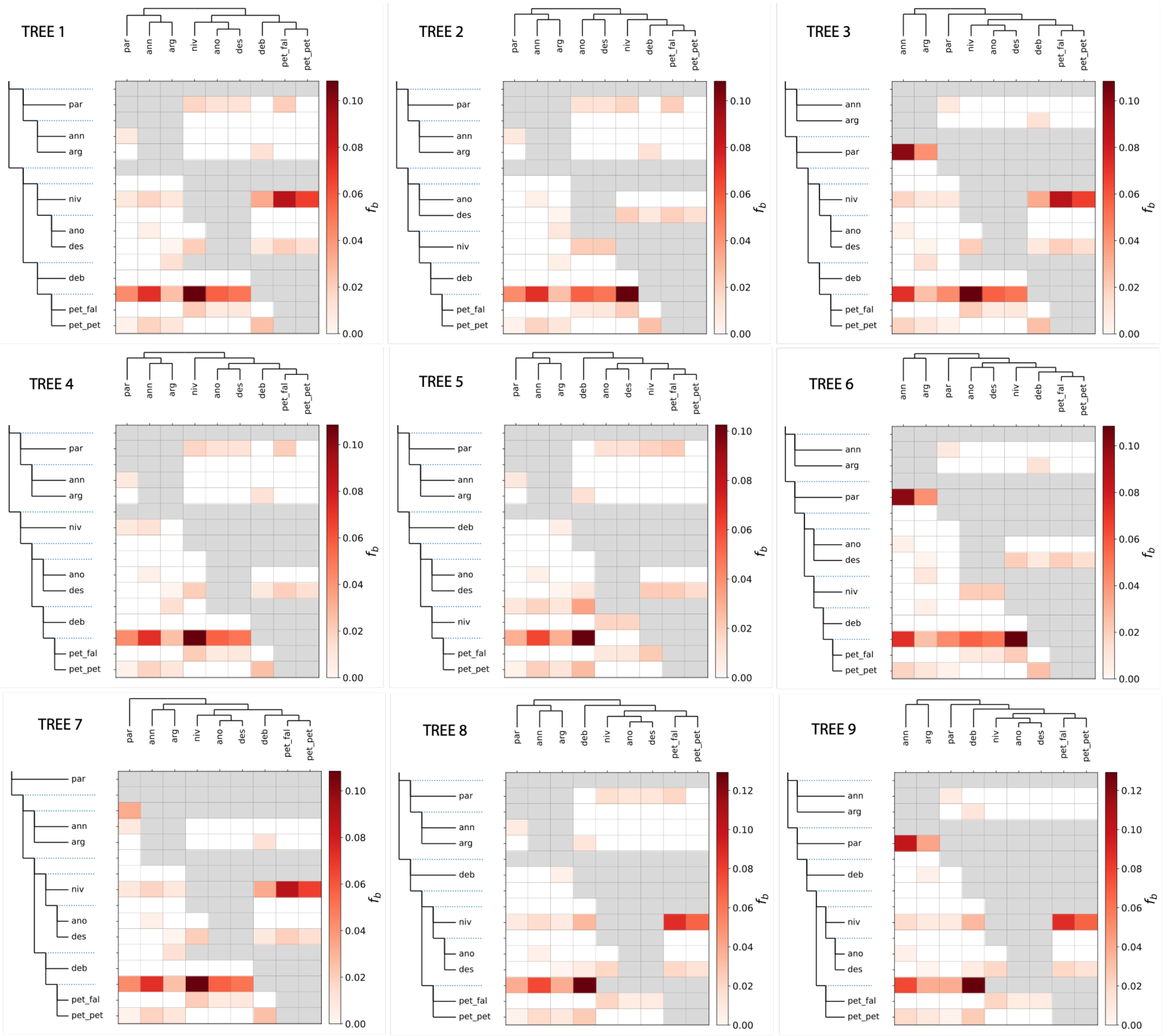


**Supplementary Figure 3: F-branch scores with alternate tree topologies.** Alternate trees were derived from the most common gene trees.

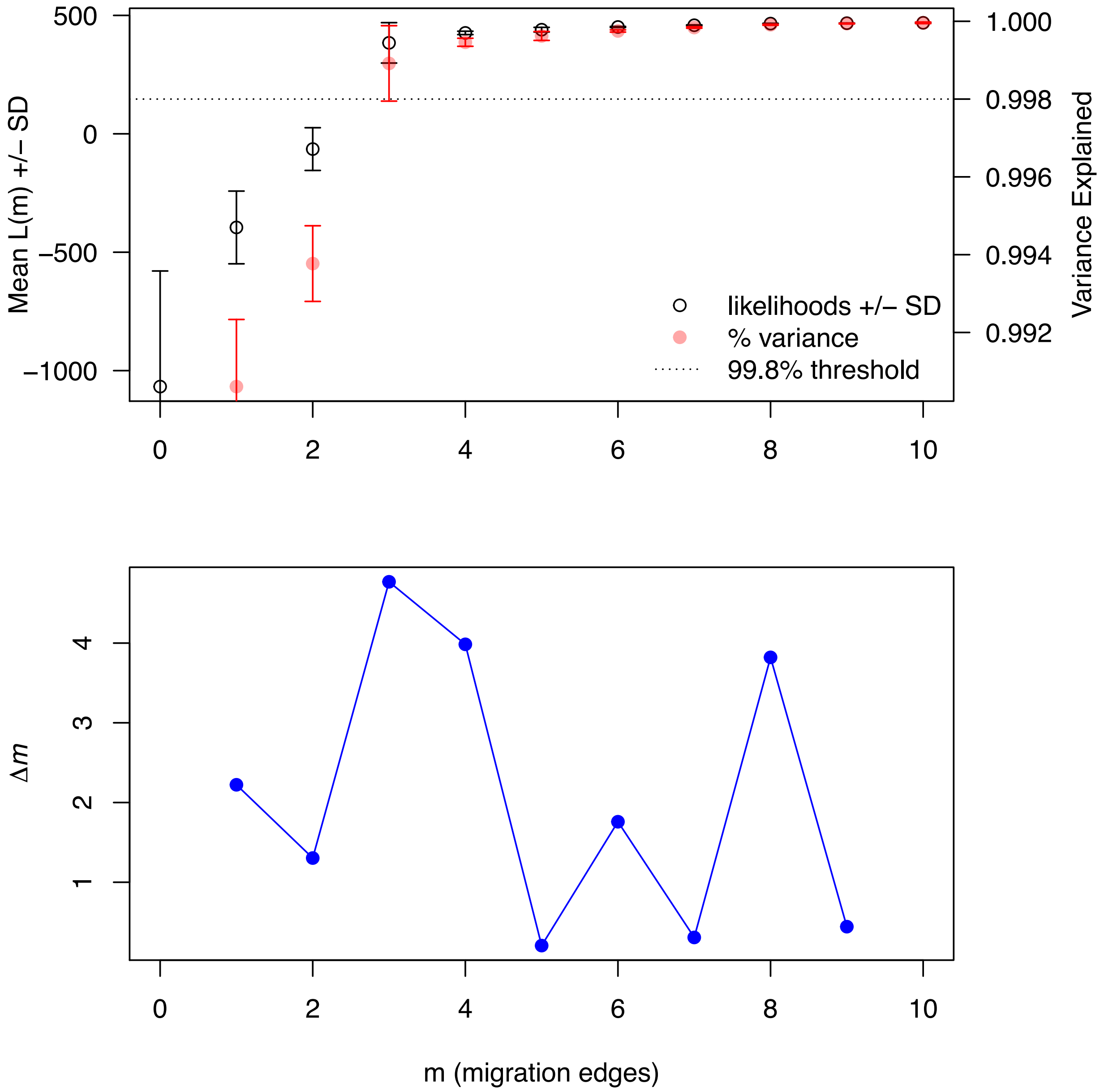


**Supplementary Figure 4: OptM chooses three as the optimal number of migration edges for TreeMix.** The mean likelihood and variance explained for 0 to 10 migration edges, as well as the second-order rate of change (Δ*m*) across values of *m*. The Evanno method was used to choose the optimal number of migration edges.

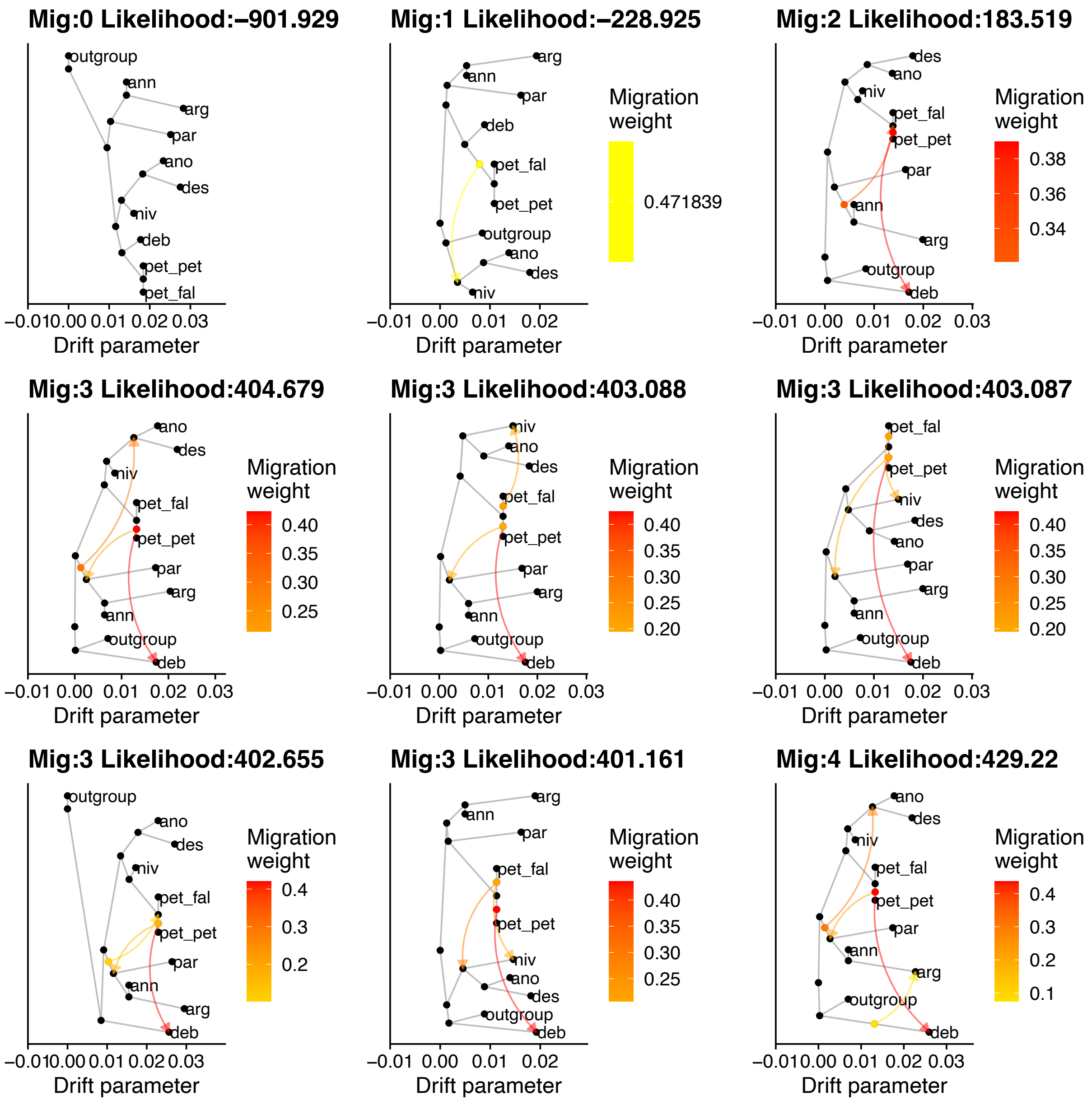

Mig:0 Likelihood:–901.929
Mig:1 Likelihood:–228.925
Mig:2 Likelihood:183.519
Mig:3 Likelihood:404.679
Mig:3 Likelihood:403.088
Mig:3 Likelihood:403.087
Mig:3 Likelihood:402.655
Mig:3 Likelihood:401.161
Mig:4 Likelihood:429.22
Migration weight
Drift parameter
outgroup
ann
arg
par
ano
des
niv
deb
pet_pet
pet_fal

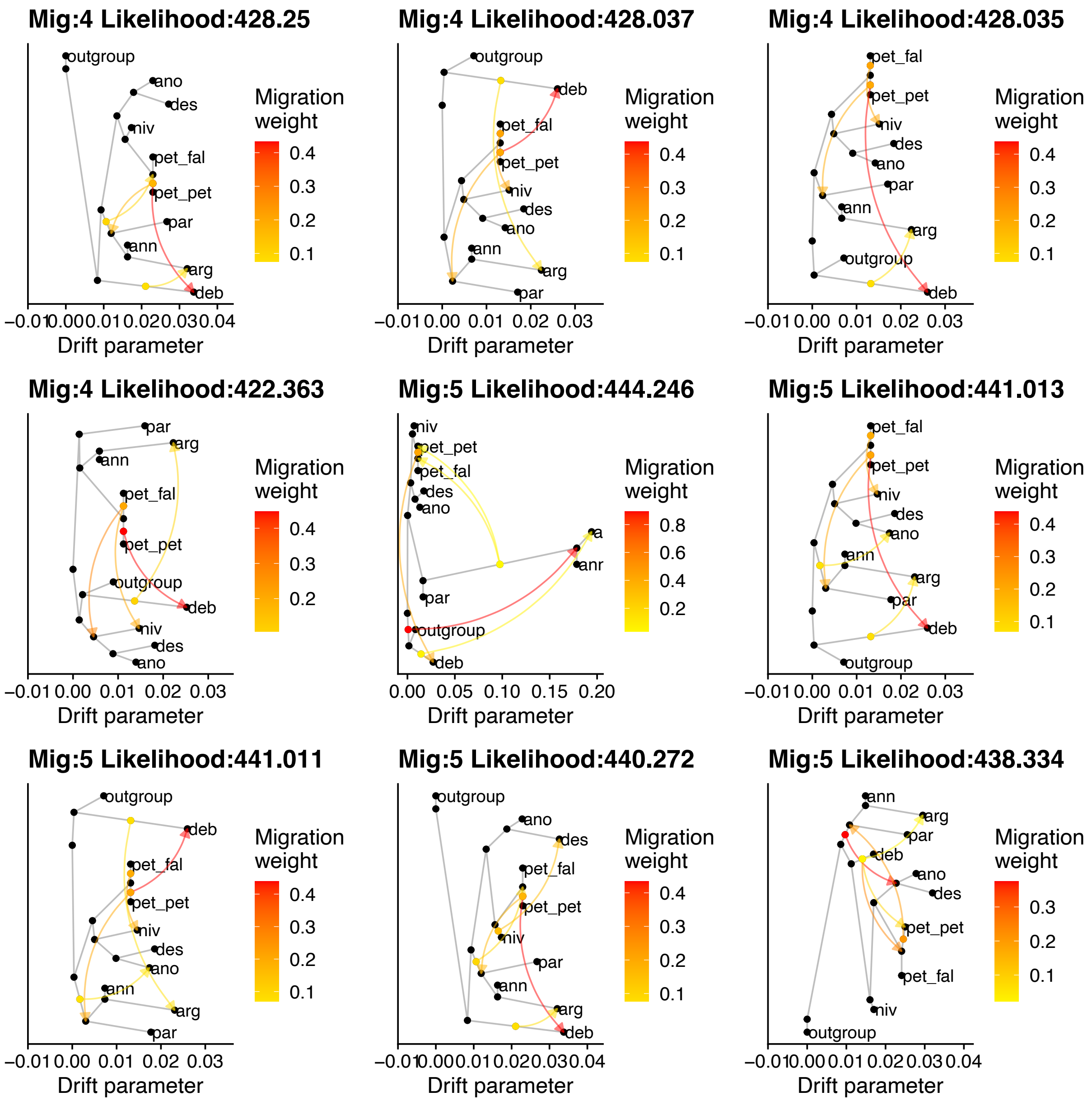
Mig:4 Likelihood:428.25
Mig:4 Likelihood:428.037
Mig:4 Likelihood:428.035
Mig:4 Likelihood:422.363
Mig:5 Likelihood:444.246
Mig:5 Likelihood:441.013
Mig:5 Likelihood:441.011
Mig:5 Likelihood:440.272
Mig:5 Likelihood:438.334
Migration weight
Drift parameter
outgroup
ano
des
niv
pet_fal
pet_pet
par
ann
arg
deb
anr

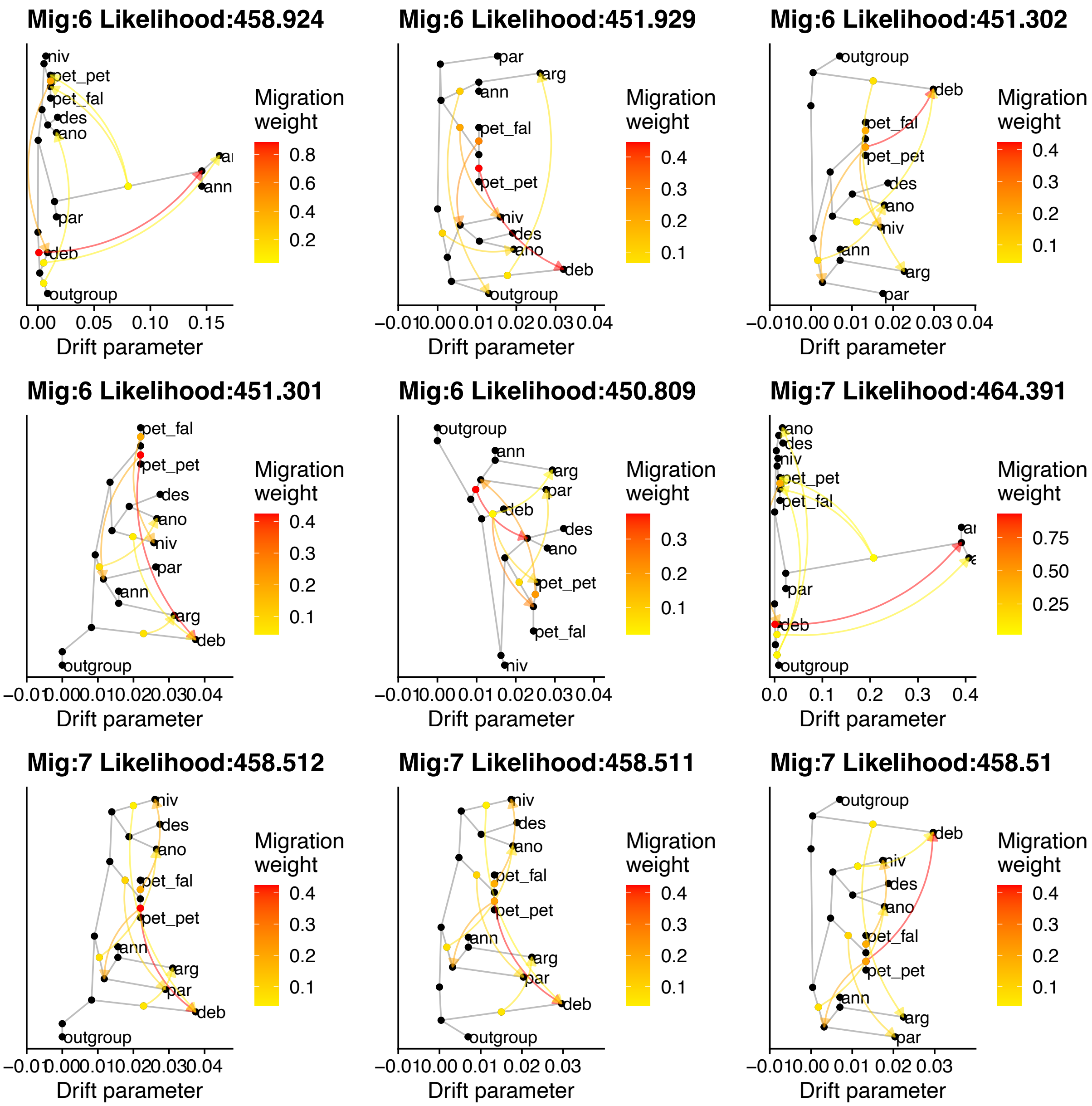

Mig:6 Likelihood:458.924
niv
pet_pet
pet_fal
des
ano
ann
par
deb
outgroup
Migration weight
0.8
0.6
0.4
0.2
0.00
0.05
0.10
0.15
Drift parameter
Mig:6 Likelihood:451.929
par
arg
ann
pet_fal
pet_pet
niv
des
ano
deb
outgroup
Migration weight
0.4
0.3
0.2
0.1
Drift parameter
Mig:6 Likelihood:451.302
outgroup
deb
pet_fal
pet_pet
des
ano
niv
ann
arg
par
Migration weight
0.4
0.3
0.2
0.1
Drift parameter
Mig:6 Likelihood:451.301
pet_fal
pet_pet
des
ano
niv
par
ann
arg
deb
outgroup
Migration weight
0.4
0.3
0.2
0.1
Drift parameter
Mig:6 Likelihood:450.809
outgroup
ann
arg
par
deb
des
ano
pet_pet
pet_fal
niv
Migration weight
0.3
0.2
0.1
Drift parameter
Mig:7 Likelihood:464.391
ano
des
niv
pet_pet
pet_fal
par
deb
outgroup
Migration weight
0.75
0.50
0.25
0.0
0.1
0.2
0.3
0.4
Drift parameter
Mig:7 Likelihood:458.512
niv
des
ano
pet_fal
pet_pet
ann
arg
par
deb
outgroup
Migration weight
0.4
0.3
0.2
0.1
Drift parameter
Mig:7 Likelihood:458.511
niv
des
ano
pet_fal
pet_pet
ann
arg
par
deb
outgroup
Migration weight
0.4
0.3
0.2
0.1
−0.01
0.00
0.01
0.02
0.03
Drift parameter
Mig:7 Likelihood:458.51
outgroup
deb
niv
des
ano
pet_fal
pet_pet
ann
arg
par
Migration weight
0.4
0.3
0.2
0.1
Drift parameter

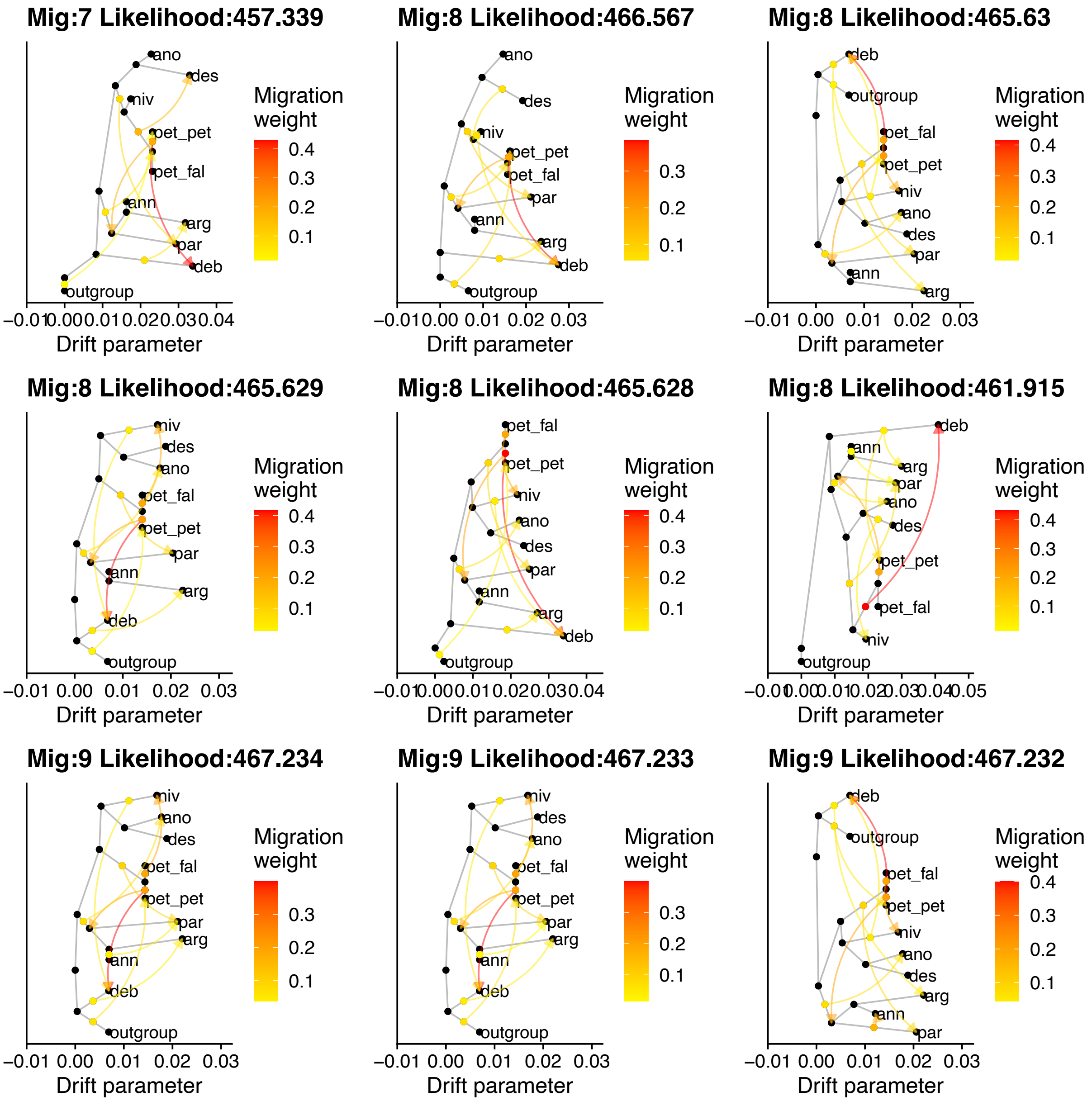


**Supplementary Figure 5: The top five TreeMix graphs within 10 likelihood of the maximum likelihood value for 0 to 10 migration nodes.**

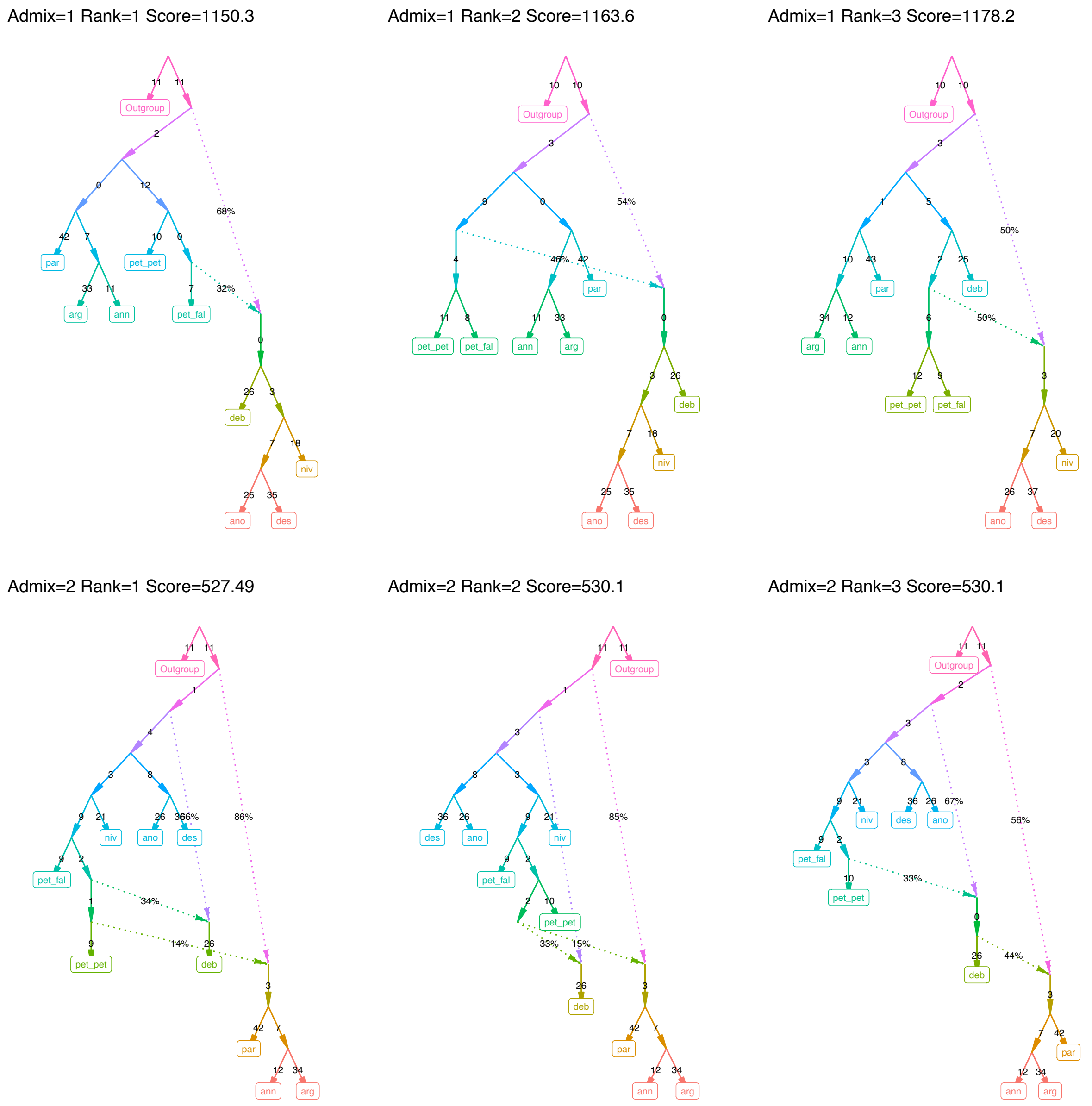

Admix=1 Rank=1 Score=1150.3
Outgroup
par
arg
ann
pet_pet
pet_fal
deb
niv
ano
des
68%
32%
Admix=1 Rank=2 Score=1163.6
Outgroup
pet_pet
pet_fal
ann
arg
par
deb
niv
ano
des
54%
46%
Admix=1 Rank=3 Score=1178.2
Outgroup
arg
ann
par
deb
pet_pet
pet_fal
niv
ano
des
50%
50%
Admix=2 Rank=1 Score=527.49
Outgroup
niv
ano
des
pet_fal
pet_pet
deb
par
ann
arg
66%
86%
34%
14%
Admix=2 Rank=2 Score=530.1
Outgroup
des
ano
niv
pet_fal
pet_pet
deb
par
ann
arg
85%
33%
15%
Admix=2 Rank=3 Score=530.1
Outgroup
niv
des
ano
pet_fal
pet_pet
deb
par
ann
arg
67%
56%
33%
44%

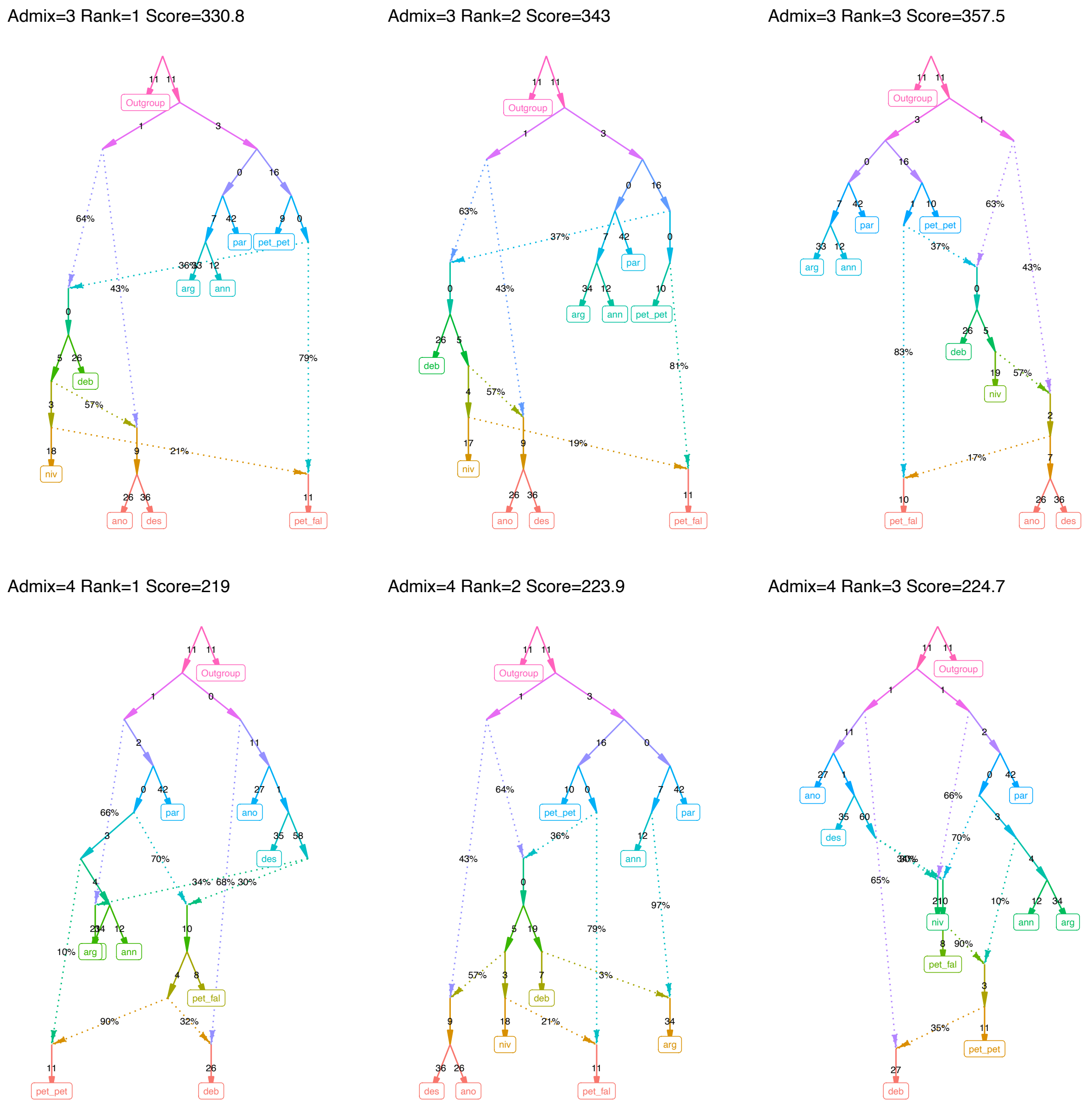


**Supplementary Figure 6: The top 3 alternate topologies for Admixtools selected admixture graphs for 1 to 4 admixture edges.** Score is the likelihood score, where smaller numbers are better.

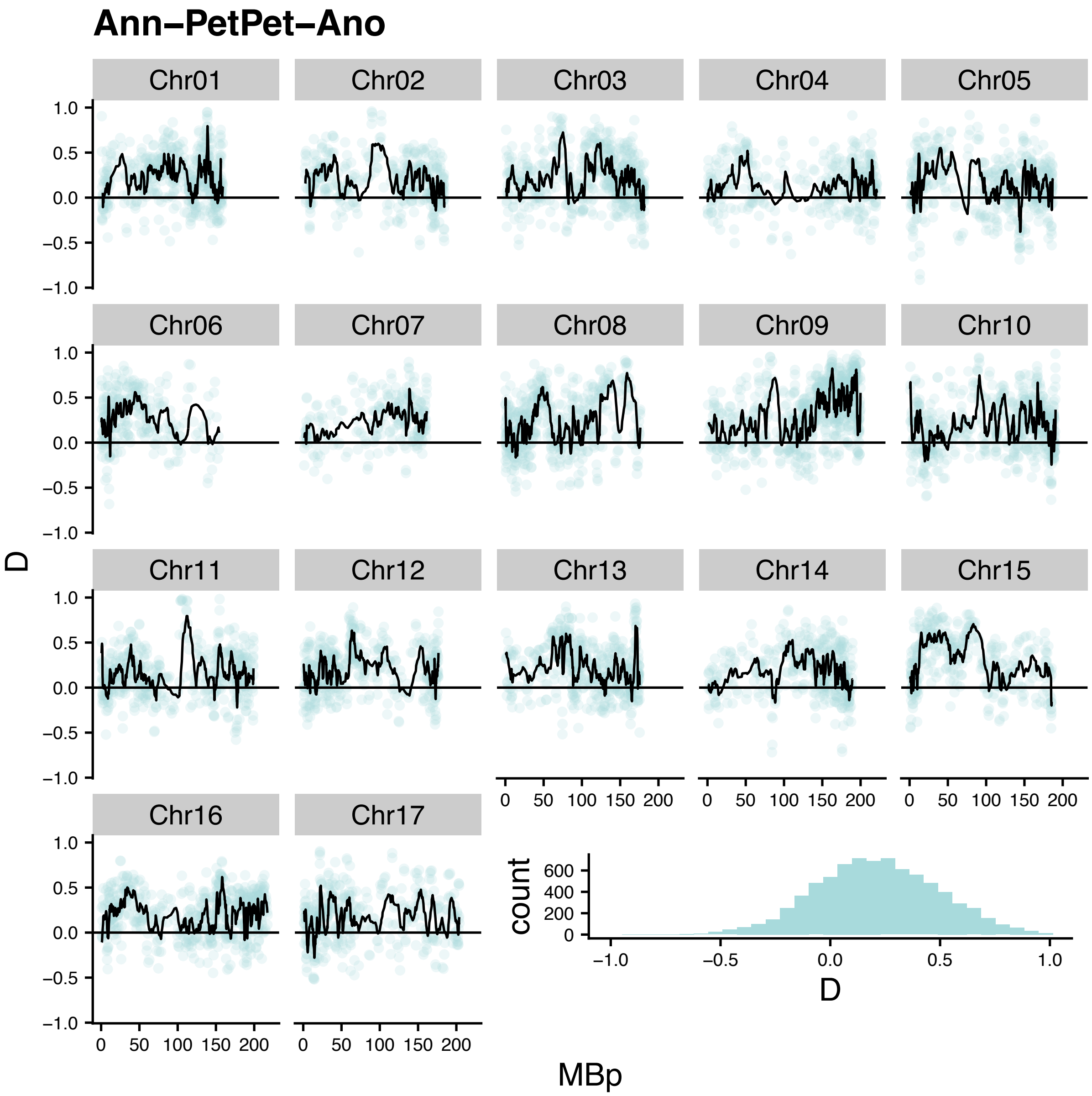
Ann−PetPet−Ano
Chr01
Chr02
Chr03
Chr04
Chr05
Chr06
Chr07
Chr08
Chr09
Chr10
Chr11
Chr12
Chr13
Chr14
Chr15
Chr16
Chr17
D
MBp
count
D

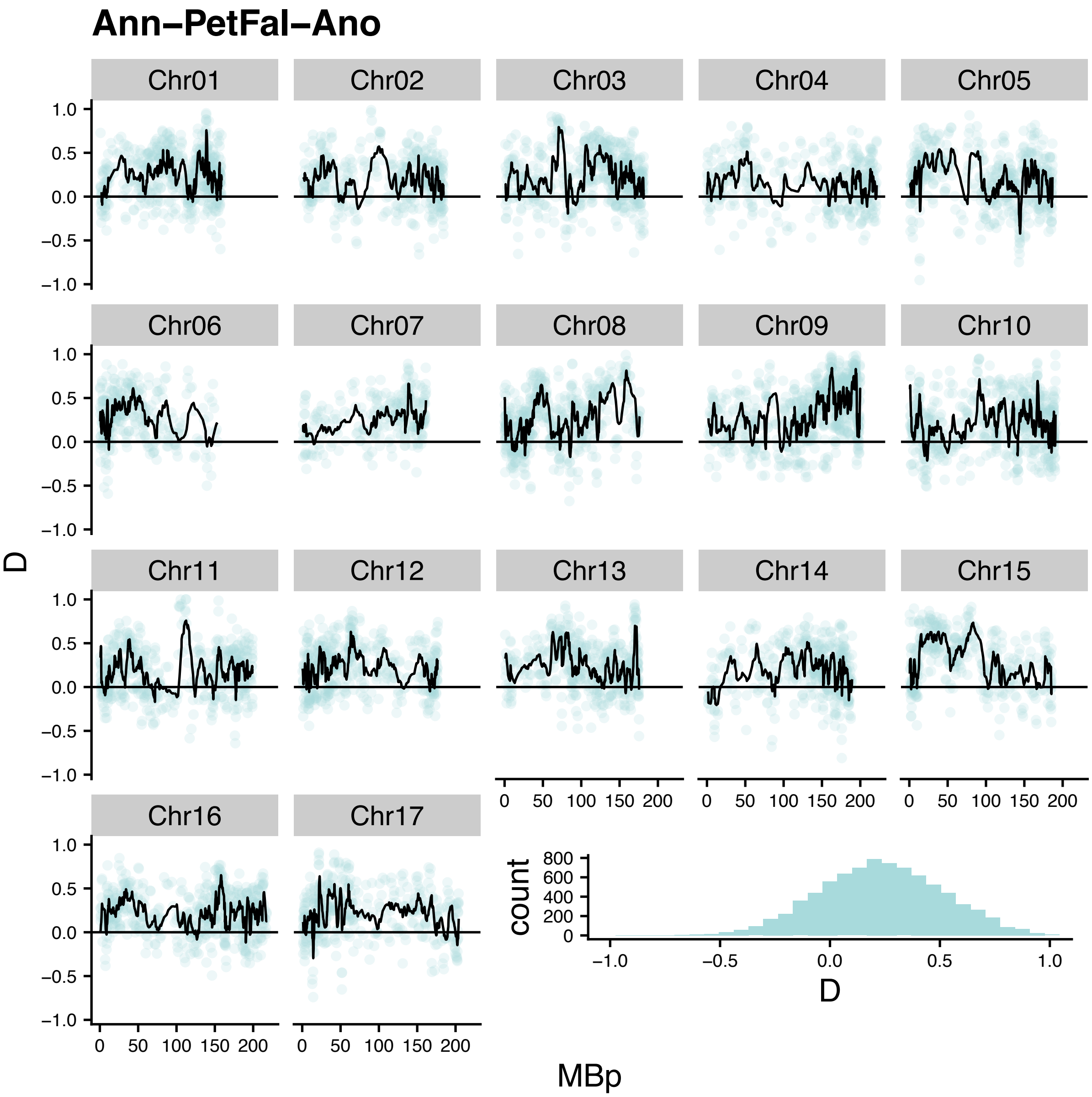

Ann–PetFal–Ano
Chr01
Chr02
Chr03
Chr04
Chr05
Chr06
Chr07
Chr08
Chr09
Chr10
Chr11
Chr12
Chr13
Chr14
Chr15
Chr16
Chr17
D
MBp
count

### Ann–Niv–Ano

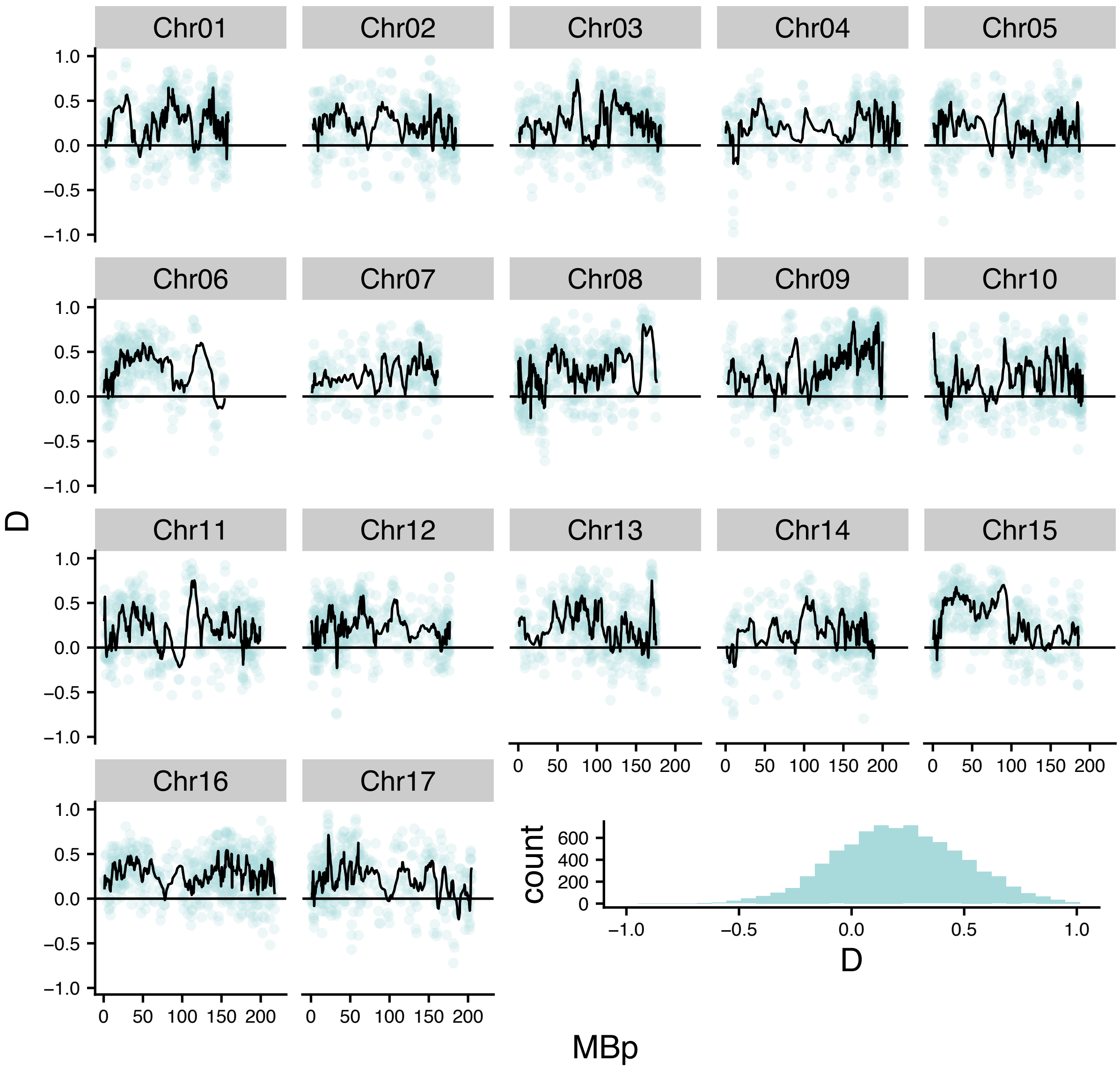

Chr01
Chr02
Chr03
Chr04
Chr05
Chr06
Chr07
Chr08
Chr09
Chr10
Chr11
Chr12
Chr13
Chr14
Chr15
Chr16
Chr17
D
MBp
count
1.0
0.5
0.0
−0.5
−1.0
0 50 100 150 200
600
400
200
0

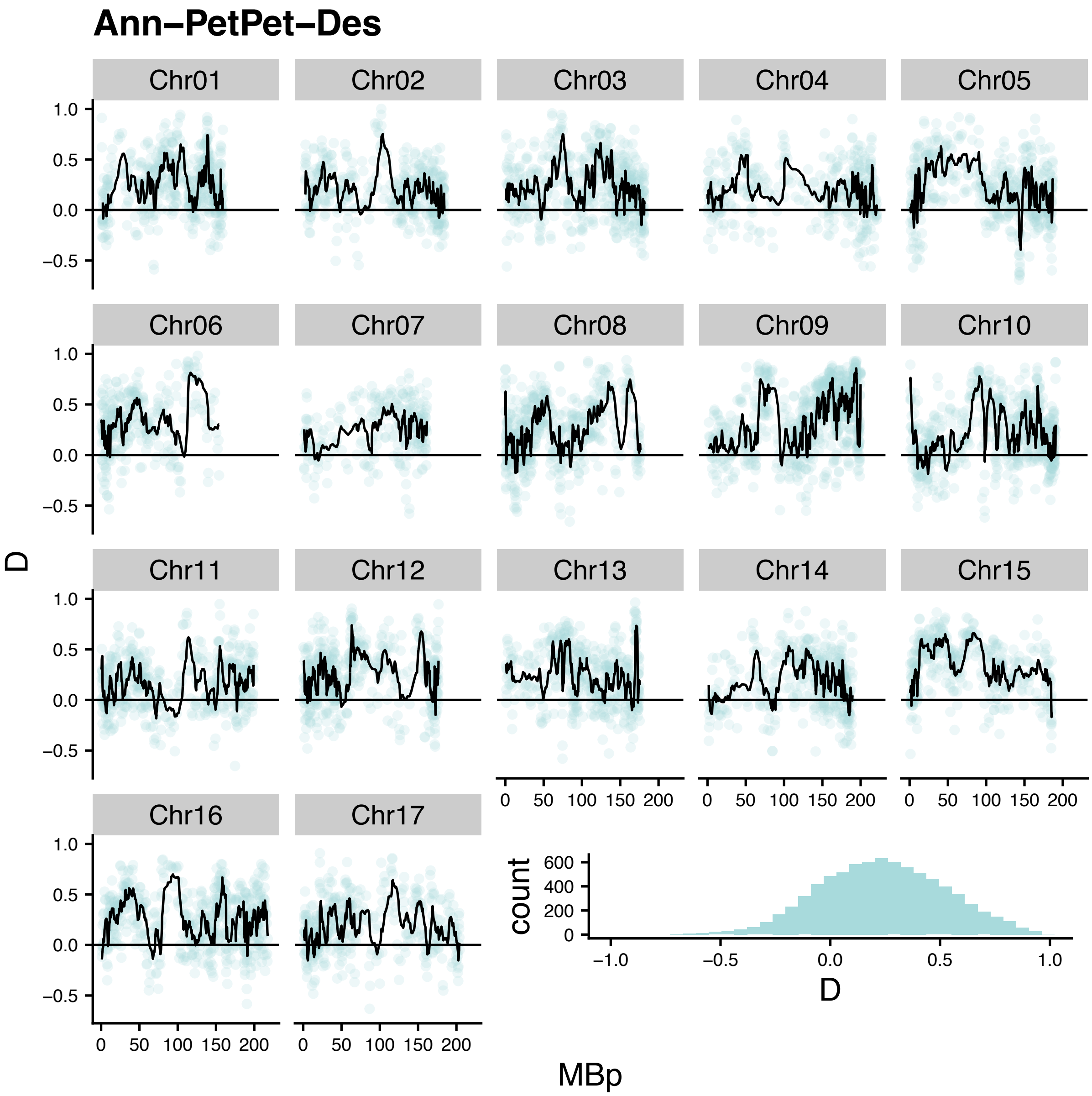

Ann–PetPet–Des
Chr01
Chr02
Chr03
Chr04
Chr05
Chr06
Chr07
Chr08
Chr09
Chr10
Chr11
Chr12
Chr13
Chr14
Chr15
Chr16
Chr17
D
count
MBp

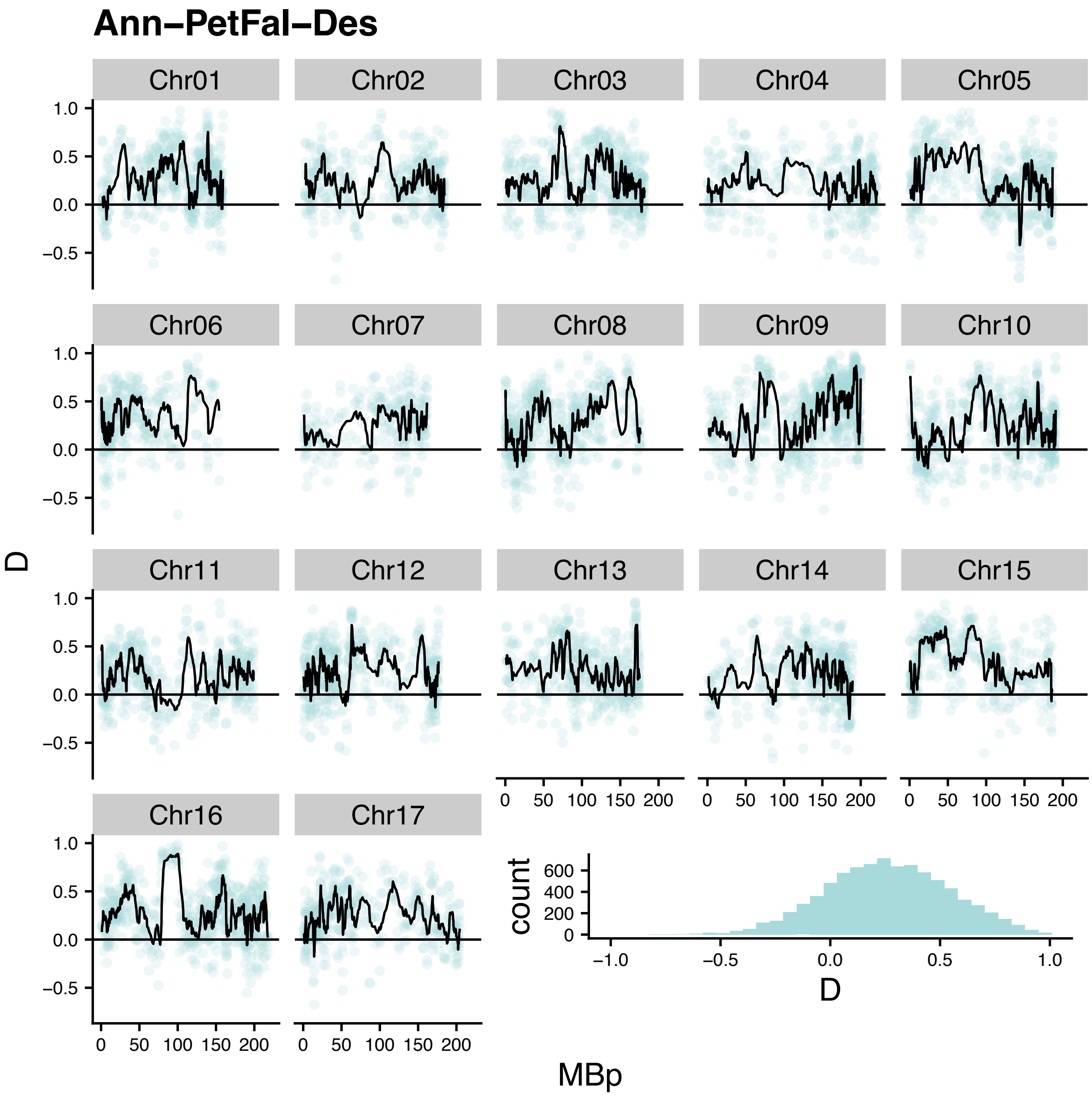

Ann–PetFal–Des
Chr01
Chr02
Chr03
Chr04
Chr05
Chr06
Chr07
Chr08
Chr09
Chr10
Chr11
Chr12
Chr13
Chr14
Chr15
Chr16
Chr17
D
count
MBp

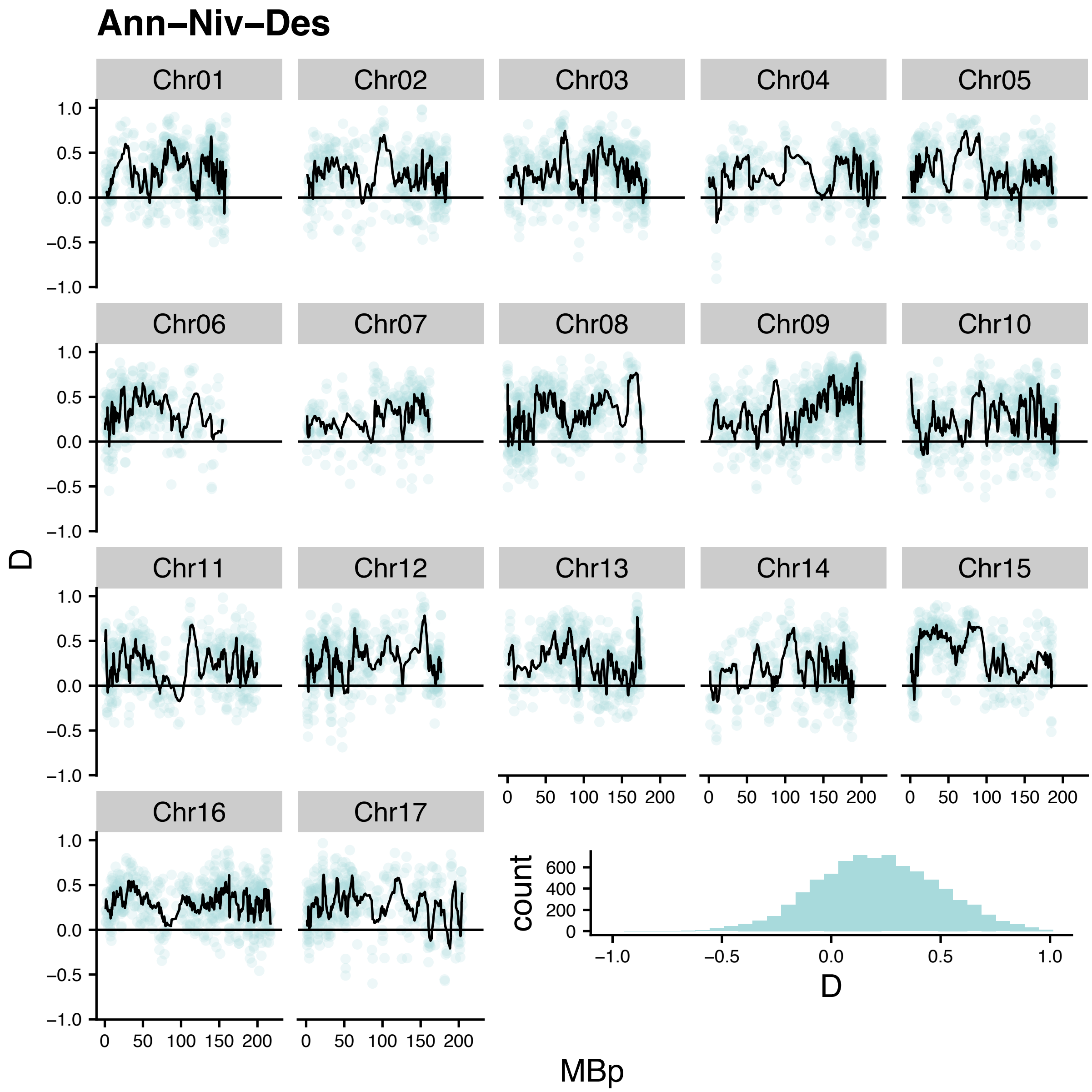

Ann–Niv–Des
Chr01
Chr02
Chr03
Chr04
Chr05
Chr06
Chr07
Chr08
Chr09
Chr10
Chr11
Chr12
Chr13
Chr14
Chr15
Chr16
Chr17
D
MBp
count
1.0
0.5
0.0
−0.5
−1.0
0
50
100
150
200
600
400
200

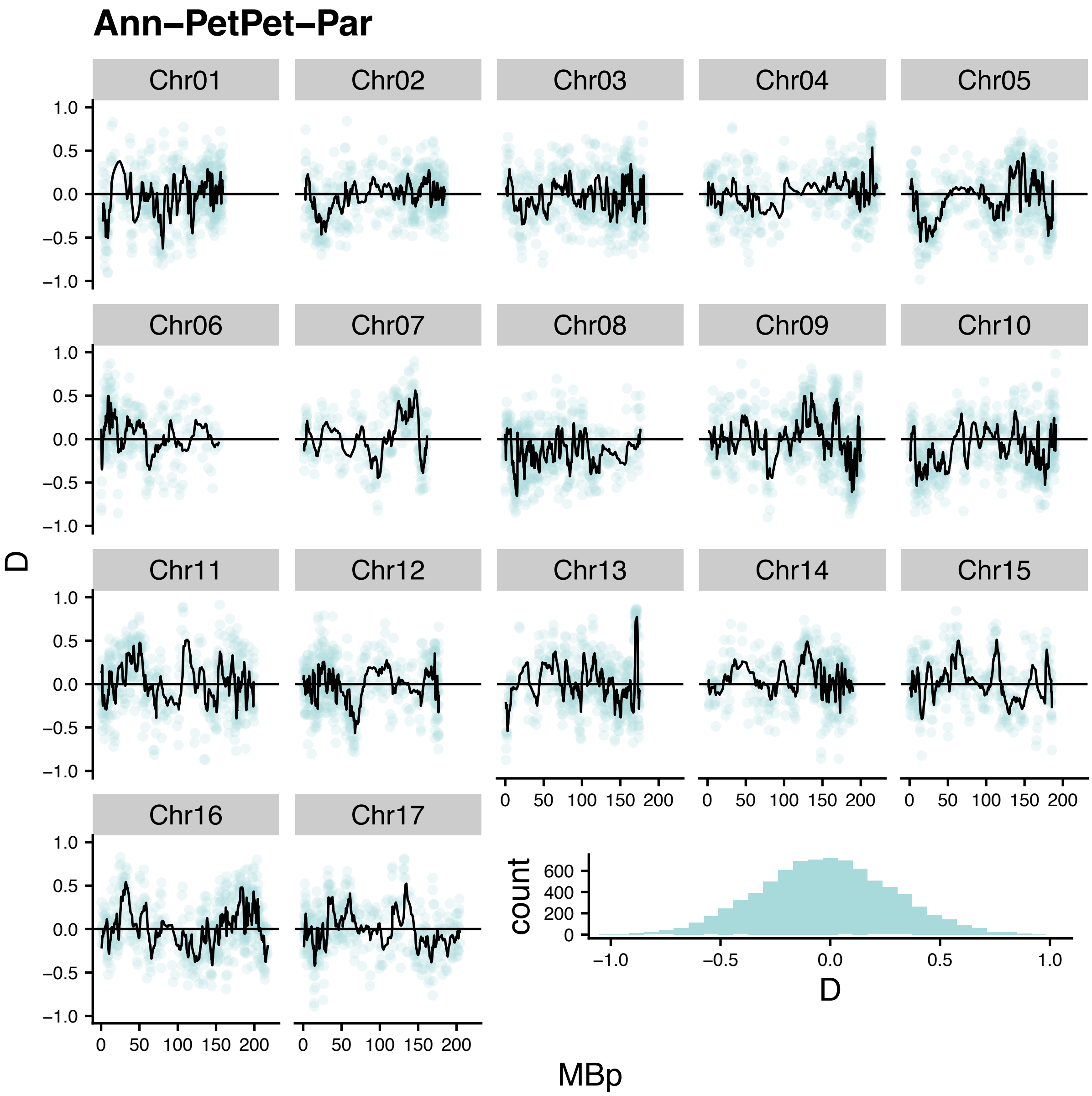
Ann–PetPet–Par
Chr01
Chr02
Chr03
Chr04
Chr05
Chr06
Chr07
Chr08
Chr09
Chr10
Chr11
Chr12
Chr13
Chr14
Chr15
Chr16
Chr17
D
1.0
0.5
0.0
–0.5
–1.0
0
50
100
150
200
MBp
count
600
400
200
0
–1.0
–0.5
0.0
0.5
1.0
D

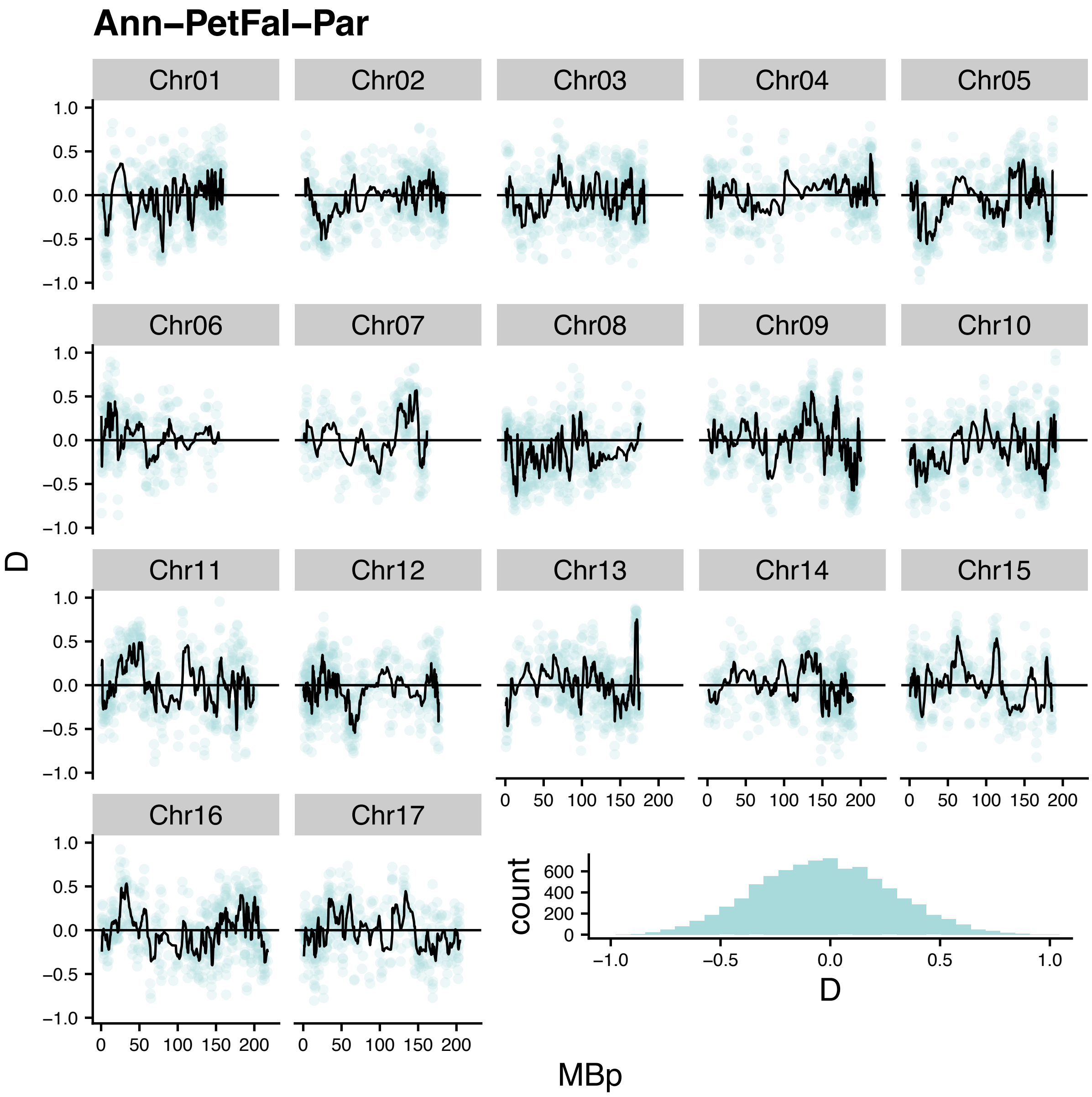
Ann−PetFal−Par
Chr01
Chr02
Chr03
Chr04
Chr05
Chr06
Chr07
Chr08
Chr09
Chr10
Chr11
Chr12
Chr13
Chr14
Chr15
Chr16
Chr17
D
MBp
count
D

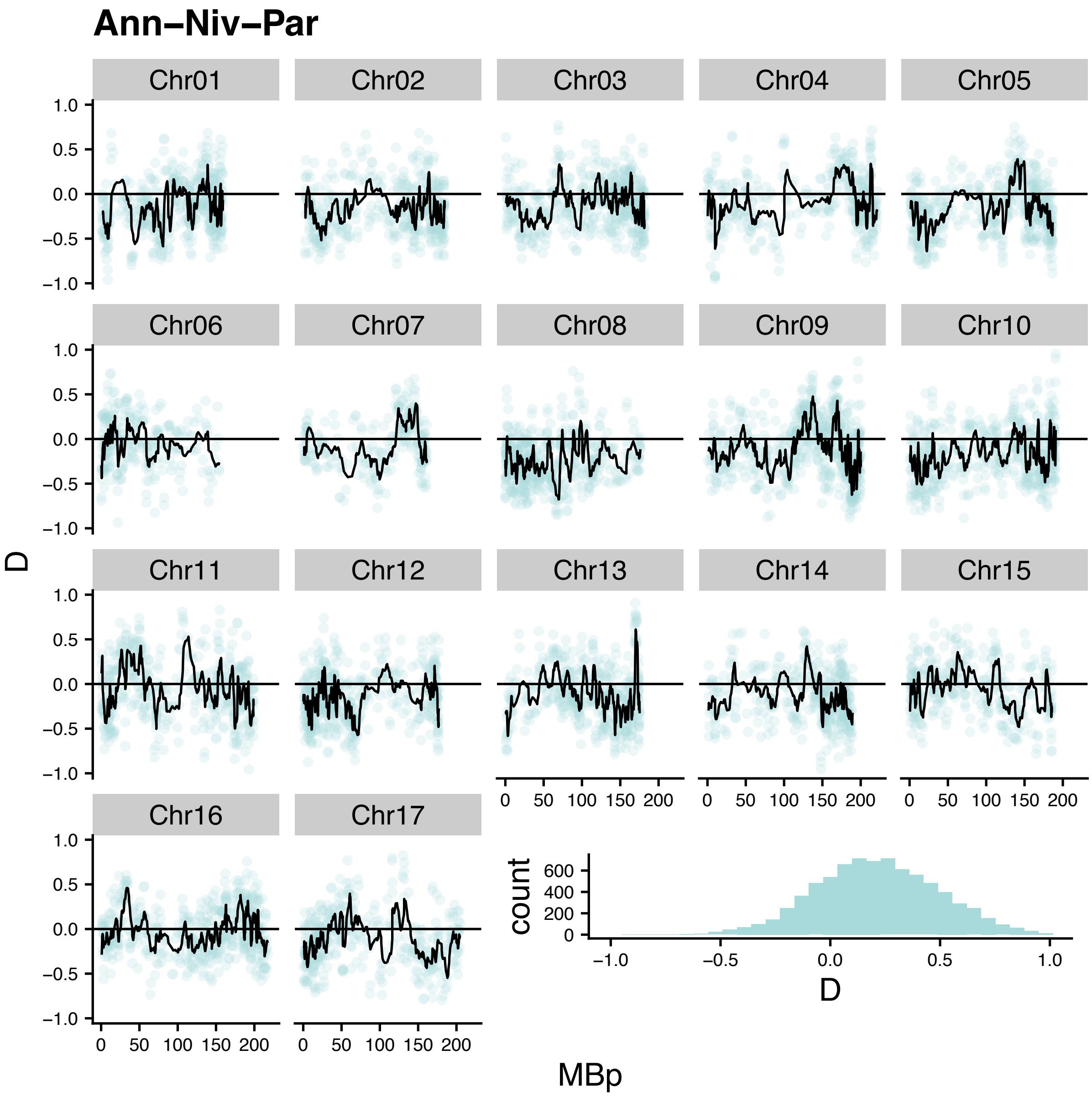


**Supplementary Figure 7: Patterson's D scores across the genome.** Values were calculated in 50 SNP windows with a 25 SNP step size. The line represents the rolling mean for 10 windows. Positive values indicate an excess of ABBA sites and negative values indicate an excess of BABA sites. The perennial species were used as an outgroup. The histogram shows the distribution of D values.

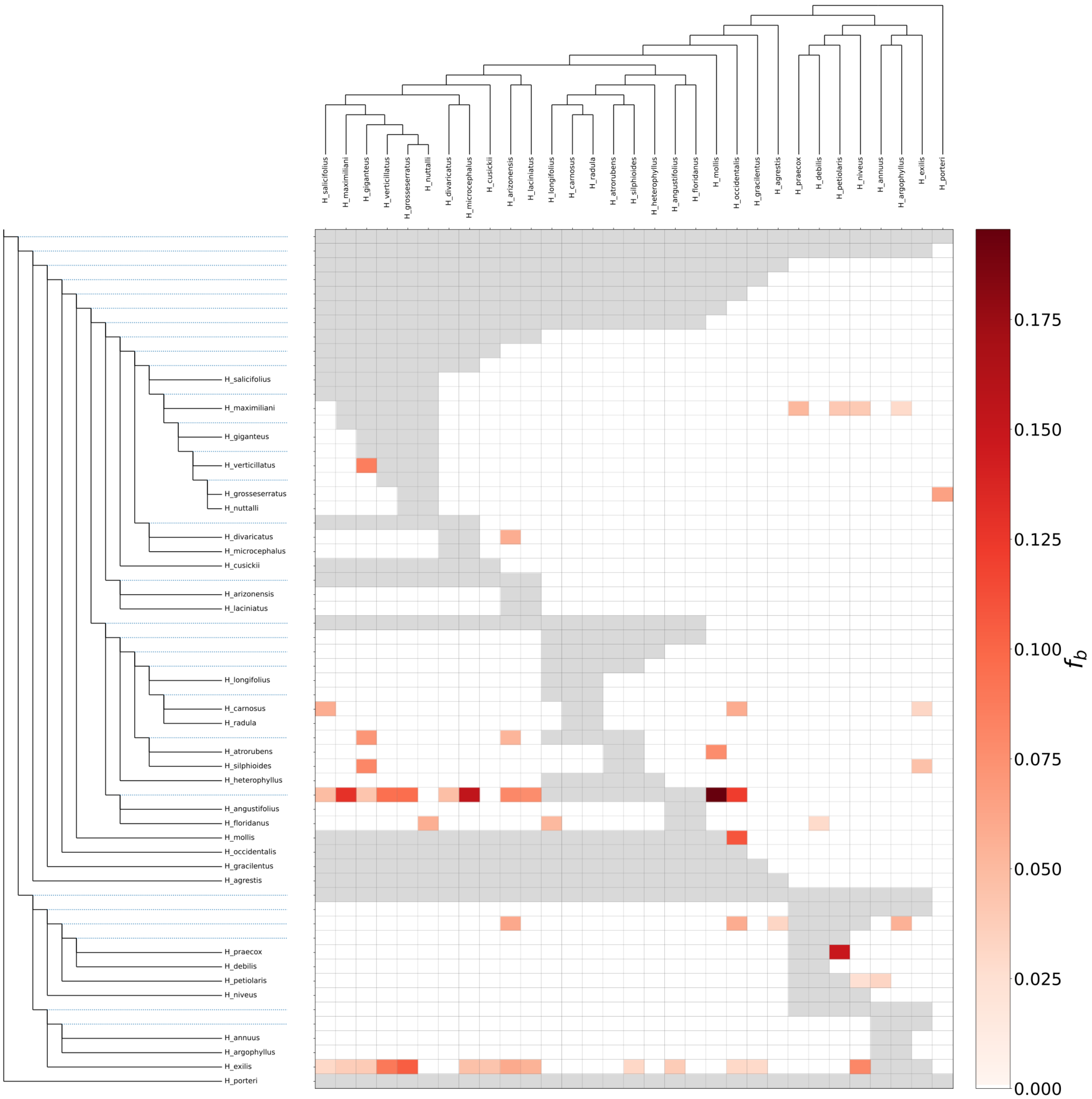


**Supplementary Figure 8: $F_{branch}$ across the *Helianthus* genus using sequence capture data.** Species tree supplied from Stephens et al. 2015. Calculations were done using Dsuite with a minimum p-value of 0.01 for inclusion.

**Supplementary Table 1: Information on samples used in analyses.**

**Supplementary Table 2: $F_{branch}$ results for all species trios in annual species dataset.**

**Supplementary Table 3: $F_{branch}$ results for all species trios in *Helianthus*-wide species dataset.**

### *Supplementary References:*


Abbott R, Albach D, Ansell S, Arntzen JW, Baird SJE, Bierne N, Boughman J, Brelsford A, Buerkle CA, Buggs R, et al. 2013 Hybridization and speciation. *J Evol Biol* 26:229-246.

Barton NH. 2001 The role of hybridization in evolution. *Mol Ecol* 2001. 10:551-568.

Barton, NH, Hewitt GM. 1985. Analysis of Hybrid Zones. *Annu Rev Ecol Evol Syst* 16:1, 113-148

Buerkle CA, Wolf DE, Rieseberg LH. 2003. The origin and extinction of species through hybridization. Pages 117-141 in C.A. Brigham, and M.W. Schwartz (Eds.), Population Viability in Plants. Springer Verlag, Berlin.

Gross BL, Schwarzbach AE, Rieseberg LH. 2003. Origin(s) of the diploid hybrid species *Helianthus deserticola* (Asteraceae). *Am J Bot* 90:1708-1719.

Lundemo S, Falahati-Anbaran M, Stenøien HK. 2009. Seed banks cause elevated generation times and effective population sizes of *Arabidopsis thaliana* in northern Europe. *Mol Ecol* 18:2798-2811.

Owens GL, Baute GJ, Rieseberg LH. 2016. Revisiting a classic case of introgression: Hybridization and gene flow in Californian sunflowers. *Mol Ecol* 25:2630-2643.

Owens GL, Samuk K. 2020 Adaptive introgression during environmental change can weaken reproductive isolation. *Nat Clim Change* 10:58-62.

Owens GL, Todesco M, Bercovich N, Légaré JS, Mitchell N, Whitney KD, Rieseberg LH. 2021. Standing variation rather than recent adaptive introgression probably underlies differentiation of the *texanus* subspecies of *Helianthus annuus*. *Mol Ecol* 30:6229-6245.

Schwarzbach AE, Rieseberg LH. 2002. Likely multiple origins of a diploid hybrid sunflower species. Mol Ecol 11:1703-1717.

Stephens JD, Rogers WL, Mason CM, Donovan LA, Malmberg RL 2015. Species tree estimation of diploid *Helianthus* (Asteraceae) using target enrichment. *Am J Bot*, 102:910-920.

Strasburg JL, Kane NC, Raduski AR, Bonin A, Michelmore R, Rieseberg LH. 2011 Effective population size is positively correlated with levels of adaptive divergence among annual sunflowers. *Mol Bol Evol* 28:1569-1580.

Todesco M, Pascual MA, Owens GL, Ostevik KL, Moyers BT, Hübner S, Heredia SM, Hahn MA, Caseys C, Bock DG, Rieseberg LH. 2016. Hybridization and extinction. *Evol Appl* 9:892-908.

Welch ME, Rieseberg LH. 2002. Habitat divergence between a homoploid hybrid sunflower species, *Helianthus paradoxus* (Asteraceae), and its progenitors. *Am J Bot* 89:472-478.

Zheng Y, Janke A. 2018. Gene flow analysis method, the D-statistic, is robust in a wide parameter space. *BMC Bioinform* 19:10.
